# Design and construction towards a pan-microbial toolkit

**DOI:** 10.1101/2024.02.23.581749

**Authors:** Charlie Gilbert, Alexander Crits-Christoph, Elise Ledieu-Dherbécourt, Shinyoung Clair Kang, Stephanie L. Brumwell, Henry H. Lee, Nili Ostrov

## Abstract

Establishing genetic tractability in non-model microbes requires identifying genetic parts that function in a target host. However, the paucity and purported narrow host range of available parts means that successful identification is governed by serendipity. Instead, a more comprehensive and scalable process would be desirable. Here, we describe the design principles for a pan-microbial genetic toolkit in which phylogenetically-diverse parts can be assembled and tested for function in microbes using high-throughput readouts. The architecture is based on Golden Gate Assembly, which simplifies the addition of parts and the construction of combinatorial libraries. We used this framework to develop two modules: first, the **POSSUM** (<u>P</u>lasmid <u>O</u>rigins and <u>S</u>election Marker<u>S</u> for <u>U</u>ndomesticated <u>M</u>icrobes) module for identification of replicating plasmids in non-model microbes which includes 29 plasmid origin of replication sequences, 23 selection markers, and 30 unique DNA sequences for tracking by sequencing; second, the **MACKEREL** (<u>M</u>odular, NGS-tr<u>ACK</u>able <u>E</u>xp<u>R</u>ession <u>EL</u>ement) module, for identification of functional gene expression cassettes which includes 426 bacterial promoter-RBS sequences driving fluorescent reporter expression, trackable by flow cytometry. We demonstrate the use of these libraries to screen for functional promoter-RBS variants in 6 non-model microbes. Continued efforts to expand this pan-microbial toolbox will accelerate efforts to improve genetic tractability and guide research across the tree of life.

## Introduction

Non-model microbes represent a vast and largely untapped resource for biotechnological applications. However, the majority of published genetic tools are generated and optimized for model organisms like *Escherichia coli* and *Saccharomyces cerevisiae*, and are rarely suitable for use in non-model microbes. The lack of characterized genetic parts, such as replicating plasmids, selection marker genes, and protein expression cassettes, necessitates laborious screening of individual constructs to identify functional tools in a new host. To enable more efficient investigation of understudied microorganisms across the tree of life, more diverse foundational genetic resources are required.

One promising avenue to achieving this is the development of widely-accessible resources that provide: (1) highly diverse, combinatorial libraries of genetic parts compatible with a broad range of microbes, and (2) validated high-throughput screening methods to rapidly identify functional genetic parts in target microbes (**Figure 1**). This design can dramatically simplify the process of identifying functional genetic parts in non-model microbes, as exemplified in several recent studies^1–8^.

**Figure 1.**
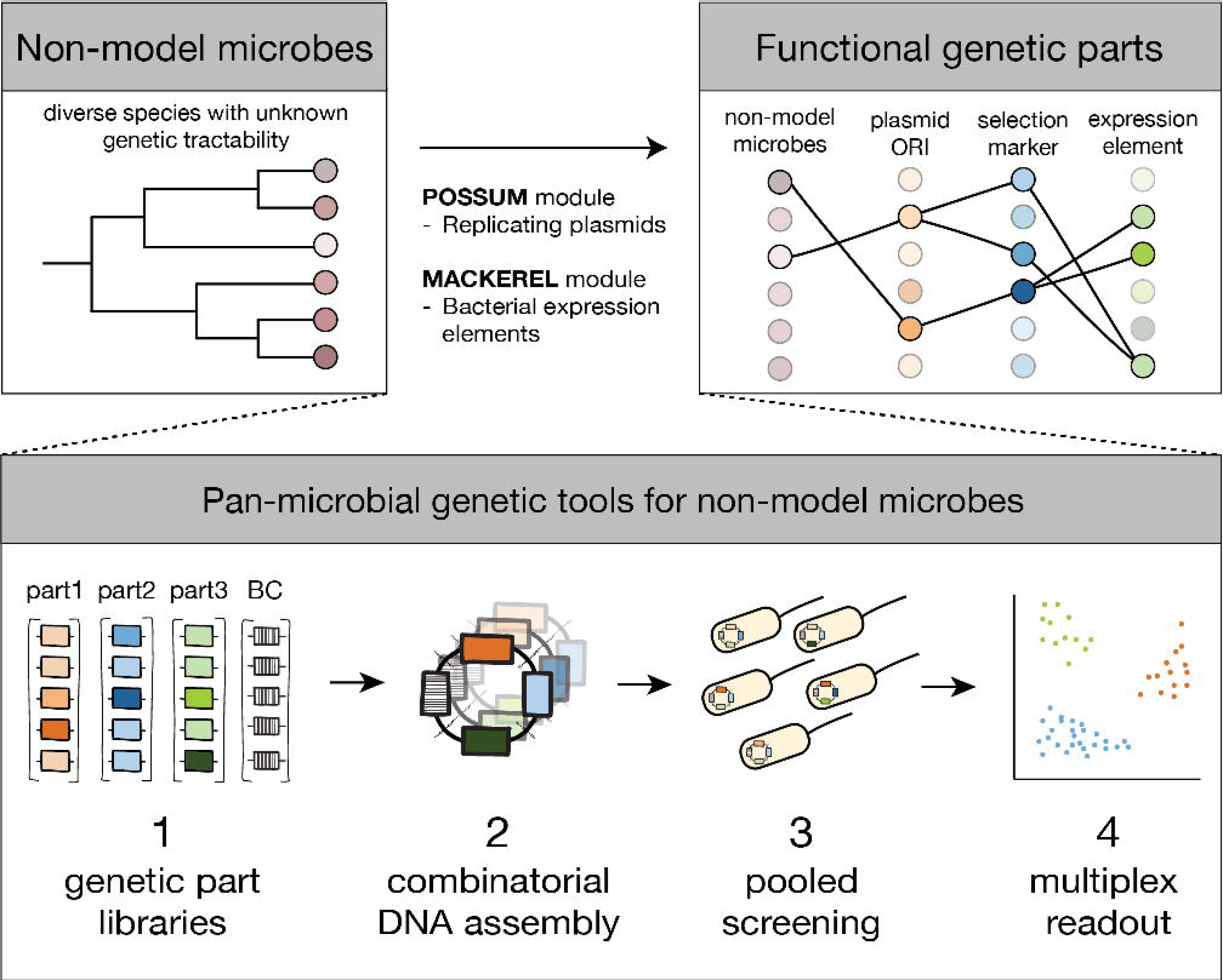
Methodological framework for expanding genetic tractability in non-model microbes. Identifying genetic parts that function in a given non-model microbe can be challenging *a priori*. One avenue to solve this challenge is to develop genetic resources consisting of: (1) libraries of barcoded (BC) genetic parts that are compatible with a broad range of hosts, (2) efficient methods for combinatorial DNA assembly, (3) high-throughput assays for functional screening of pooled libraries in target microbes, and (4) multiplexed readout analysis for identification of functional library members.

While a number of existing genetic toolkits have been developed for use in multiple microbes (**Supplementary Table 1**)^9–18^, none have been designed specifically to support scalable screening of functional genetic parts in non-model bacteria. The Standard European Vector Architecture (SEVA) collection is generally considered the largest collection of genetic parts applicable for a broad range of microbes^10–13^. However, adding or swapping parts with this collection relies on cumbersome restriction-ligation methods which have been superseded over the last decade by more efficient Golden Gate and homology-based cloning methods^19–21^. More recently a subset of SEVA parts have been modernized to fit Golden Gate assembly standards^14,18,22^, while others have introduced novel toolkits including broad host range parts^15–17^. However, overall, no single system seeks to provide a large number of genetic parts that can be assembled and tested with high multiplexability for diverse non-model microbes.

In this manuscript, as part of our work towards a pan-microbial toolkit, we describe the construction of two modules for establishing genetic tractability in non-model microbes. The **POSSUM** (<u>P</u>lasmid <u>O</u>rigins and <u>S</u>election Marker<u>s</u> for <u>U</u>ndomesticated <u>M</u>icrobes) module is a Golden Gate assembly toolkit enabling the identification of replicating plasmids in target microbes. The **MACKEREL** (<u>M</u>odular, NGS-tr<u>ACK</u>able <u>E</u>xp<u>R</u>ession <u>EL</u>ement) module is a Golden Gate-compatible, broad host range expression element pooled plasmid library, consisting of 426 bacterial promoter-RBS sequences, developed by Yim et al.^23^, which can be screened for function in target microbes. We demonstrate the utility of this approach by applying this toolbox to 6 non-model microbes in pooled screens.

Several features make both resources especially useful for new organisms and offer a unified approach for genetic tool discovery: First, highly diverse collections increase the chance of identifying functional genetic parts even in broad taxonomic groups. Second, the systems described are designed to be compatible with high-throughput screening for parallel assessment of part functionality, including DNA barcodes for pooled NGS assays exploiting flow cytometry and growth-dependent selection. Finally, by adopting a widely-used Golden Gate assembly framework, we maximize the ability to easily add new parts by synthesis and through cross-compatibility with existing genetic toolkits.

## Results and Discussion

### POSSUM module design and composition

The POSSUM module provides: (1) a collection of 126 modular genetic parts that are needed to support plasmid replication in diverse microbes, and (2) a panel of Golden Gate assembly destination vectors enabling combinatorial assembly of modular parts (**Figure 2**). Individual genetic parts are formatted as “Level 0” plasmids, in which functional parts are hosted on a plasmid backbone for ease of storage and propagation. Level 0 part types include a set of plasmid origin of replication sequences (ORIs), selection marker components and a series of additional sequences useful for engineering a range of bacterial and fungal species. The POSSUM module adheres to an existing Golden Gate toolkit architecture: the Universal Loop Assembly (uLoop)^15^ system (**Supplementary Figure 1**) which adopts the widely-used “common syntax” for Golden Gate assembly^24–26^. As a result, POSSUM module parts are cross-compatible with numerous existing Golden Gate toolkits, including JUMP^14^, MoClo^27^, GoldenBraid^28^ and others (**Supplementary Table 2**).

**Figure 2.**
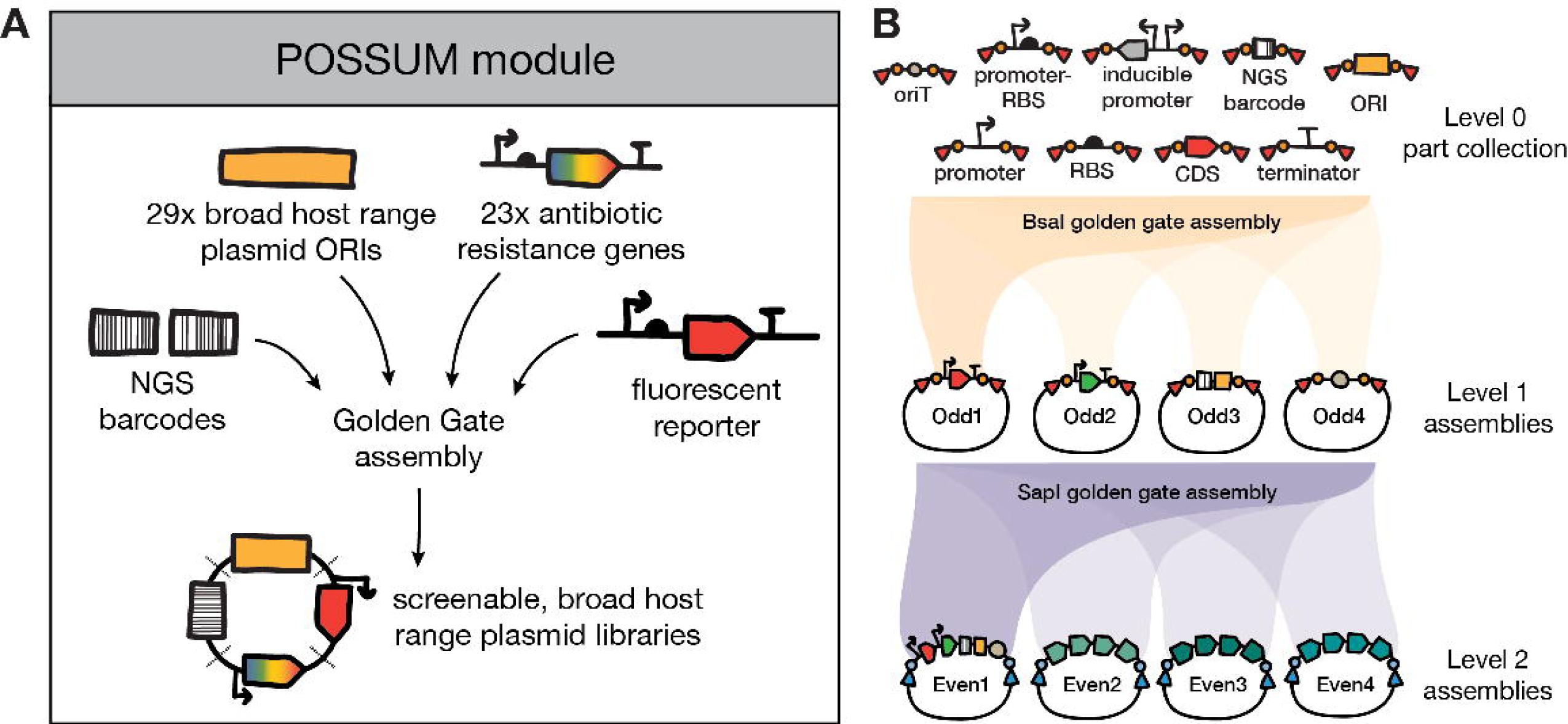
POSSUM module overview. (A). The POSSUM module provides a collection of DNA sequences which can be combined through Golden Gate assembly to create libraries of plasmid variants that can be screened for function in target microbes. (B) Illustration of the Loop Assembly method employed by the POSSUM module. Alternating rounds of BsaI and SapI Golden Gate assemblies combine up to four parts at a time into destination vectors. Assembly levels alternate between Odd and Even destination vectors. A detailed description of all POSSUM module Level 0 parts is provided in **Supplementary Table 4**. *oriT*: conjugation origin of transfer; RBS: ribosome binding site; CDS: coding sequence; NGS: next-generation sequencing.

We first generated a collection of 22 assembly destination vectors following the uLoop design (**Supplementary Table 3** and **Supplementary Figure 2A-D**). Under this design, users begin by assembling Level 0 parts into one of four Odd-level destination vector positions (Odd1-4). The parts contained in the four Odd-level destination vector positions can then be combined into one of four Even-level destination vector positions (Even1-4). This procedure can be further iterated to generate larger and larger constructs, oscillating between Odd- and Even-levels and combining components in sets of up to four at a time (**Figure 2B**). We created two versions of each assembly destination vector, differing in the *E. coli* antibiotic resistance gene used (kanamycin/carbenicillin for Odd-level, chloramphenicol/carbenicillin for Even-level). In addition, we generated six non-standard destination vectors for specific use cases such as storage of Level 0 parts and creating suicide plasmids (**Supplementary Figure 2D**).

Next, we created a collection of 126 Level 0 parts to provide the components required to screen for functional plasmids in non-model bacteria and fungi (**Supplementary Figure 3** and **Supplementary Table 4**). Level 0 parts are the fundamental building blocks of a Golden Gate toolkit and typically consist of minimal functional genetic units, e.g. promoters, protein coding-sequences (CDSs) and terminators. All Level 0 parts underwent the same DNA sequence ‘domestication’ process in which internal BsaI/SapI cut sites were removed and assembly schema added according to the Golden Gate assembly common syntax^26^.

We selected 36 ORIs known to function in a variety of bacteria and fungi, sourced from published descriptions of broad host range plasmids and genetic toolkits developed for diverse microbes. Level 0 ORI parts were generated by commercial gene synthesis or PCR amplification from existing plasmids. We successfully built 29 of 36 ORIs as Level 0 parts, while the remaining seven ORIs failed at a variety of stages in the part construction process (pVS1, pBAV1K-T5, pLF1311, pLS1, pCM62, pAP1, pWV01) (**Supplementary Table 5**). For ORIs that we were not able to construct, we did not determine the specific cause of failure. One possibility is that certain ORIs cannot be hosted on the same plasmid as the ColE1 ORI; however, the pCM62 ORI is derived from the RK2 ORI which was successfully created as a Level 0 part here and previous reports have described successful assembly of pVS1 and pLS1 ORIs with the ColE1 ORI on a single plasmid^16,29^. Finally, we created a non-functional “dummy ORI’’ control part composed of a randomly-generated 48 bp DNA sequence.

The majority of methods used to establish plasmid transformation rely on incorporation of a functional selection marker on delivered plasmid DNA, most commonly an antibiotic resistance gene. Since microbes differ in both their native antibiotic susceptibility and in the specific resistance genes that can function in them, we sourced a collection of 23 diverse antibiotic resistance genes. Each gene was split into component promoter, CDS and terminator Level 0 parts to enable modular engineering of selection markers (**Supplementary Table 6**).

To facilitate multiplexed screening of plasmid variants, we generated a set of 30 NGS barcodes as Level 0 parts. These barcode sequences are designed to allow simple and cost-effective identification of ORI variants through amplicon-sequencing of a short sequence which is uniquely paired to each ORI during DNA assembly. Barcodes consist of a variable 20 bp sequence flanked by constant 30 bp sequences which act as universal primer binding sites for amplicon-sequencing. For variable barcode regions we used the “assembly subpool-specific primers” presented in the DropSynth method, which were designed for optimal performance in PCR: lacking sequence homology, secondary structure, primer dimers and self-dimers^30,31^.

Finally, a set of additional Level 0 parts were generated to facilitate genetic manipulation of target microbes (**Supplementary Table 4**). This includes a collection of seven constitutive and three inducible promoters, four fluorescent reporter genes, six transcriptional terminators and an origin of transfer (*oriT*) sequence for the commonly-used RP4/RK2 conjugation machinery^32^.

### Using the POSSUM module to assemble ORI-marker libraries

We next used the POSSUM module to assemble combinatorial libraries of plasmids with variable ORIs and selection markers which we refer to as ORI-marker libraries (**Figure 3A**). The intended purpose of ORI-marker libraries is to provide researchers with a single, comprehensive resource to identify replicating plasmids in target microbes. To this end, we have previously described an experimental screen to identify functional plasmids from ORI-marker libraries in non-model microbes through pooled library screening^8^.

**Figure 3.**
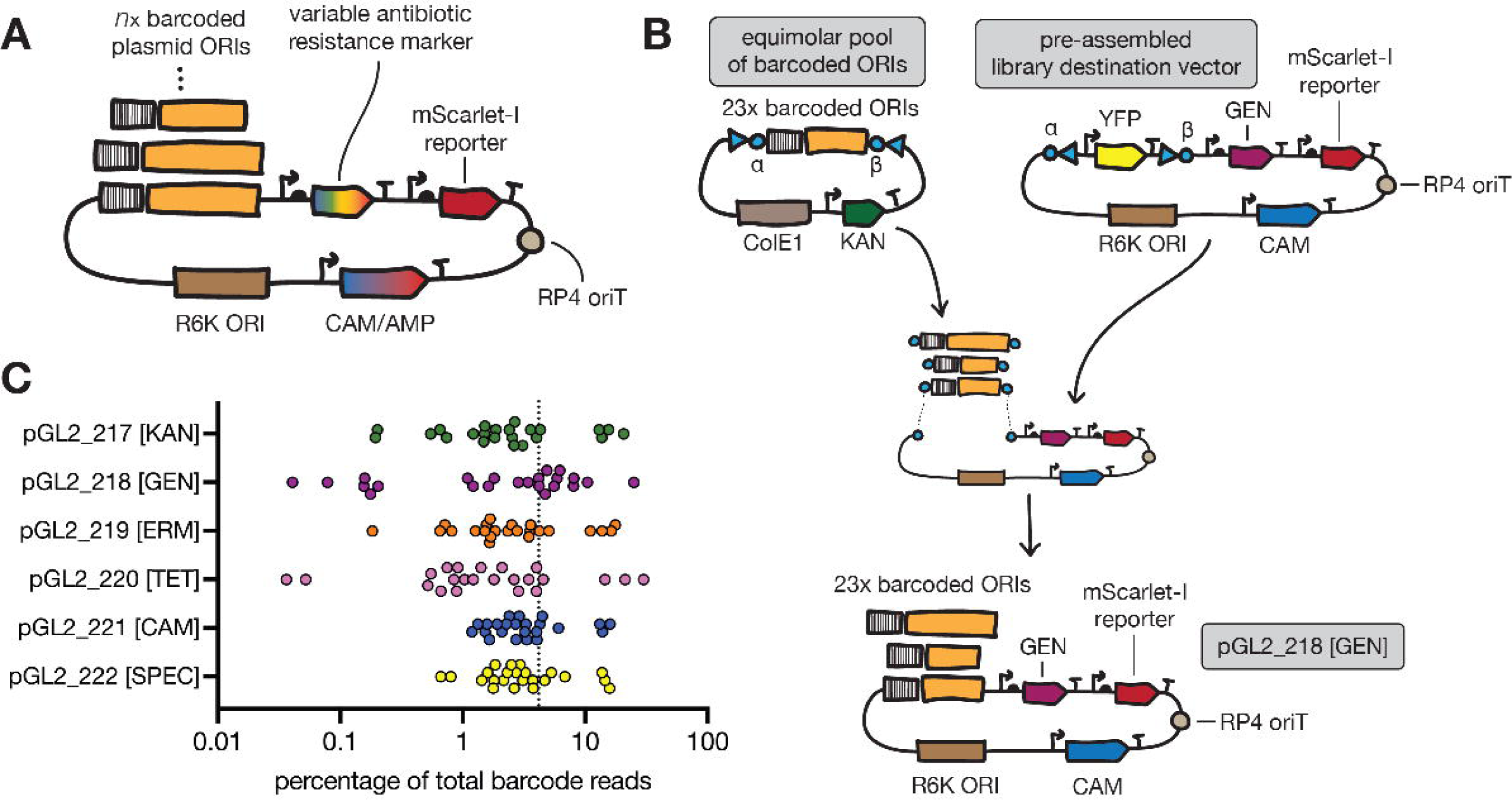
Assembly of ORI-marker libraries using the POSSUM module. (A) Schematic showing the design of ORI-marker plasmids. (B) Schematic illustrating the final Golden Gate assembly step used for constructing ORI-marker plasmid libraries. An equimolar pool of barcoded ORI Level 1 plasmids is combined with a pre-assembled destination vector carrying a selection marker (in this case gentamicin, GEN). A SapI Golden Gate assembly reaction replaces the YFP dropout part with the pooled barcoded ORIs. Assembly products are electroporated directly into an *E. coli* conjugation donor strain to create an input library. (C) Amplicon-sequencing data showing the abundance of the 23 individual ORIs within six ORI-marker libraries (pGL2_217-222). Each ORI library carries a different antibiotic selection marker (Y-axis). Amplicon-sequencing was performed on plasmid DNA extracted from the conjugation donor (‘input library’). The scatter plot illustrates the percentage of reads mapping to each ORI barcode within the libraries. The vertical dotted line represents the expected abundance of each ORI if library distribution was perfectly even.

In brief, ORI-marker plasmids are made up of: a conditional *E. coli* ORI (R6Kγ), an *E. coli* selection marker (CAM/AMP), the RP4/RK2 conjugation machinery *oriT* sequence, an mScarlet-I reporter gene, an NGS-barcoded plasmid ORI and an antibiotic resistance gene for selection in target microbes. Here, we describe the use of the POSSUM module for construction of six ORI-marker libraries, each consisting of 22x bacterial ORIs and one of six different selection markers (KAN, GEN, ERM, TET, CAM or SPEC) known to function across a range of bacterial phyla (**Supplementary Table 7**).

ORI-marker libraries were constructed from Level 0 parts in three stages. First, Level 0 parts were assembled to create four types of Level 1 constructs: (1) 22 plasmid ORIs and one dummy ORI each paired with a unique NGS barcode; (2) six antibiotic resistance cassettes; (3) a single mScarlet-I reporter expression cassette; (4) a single *oriT* cassette (**Supplementary Figure 4A-D** and **Supplementary Table 8**). Second, the Level 1 resistance cassette, reporter cassette and oriT were assembled with a YFP dropout part, resulting in six “pre-assembled library destination vectors” (**Supplementary Figure 5** and **Supplementary Table 8**). In the third and final assembly step, depicted in **Figure 3B**, 23 Level 1 barcoded-ORI plasmids were pooled and assembled into the pre-assembled library destination vectors, replacing the YFP dropout part (**Supplementary Table 8**). Final library assembly products were electroporated directly into a conjugation donor strain (*E. coli* BW29427) to create ORI-marker libraries.

While the second and third steps above could be condensed into a single assembly step, as Golden Gate assembly efficiency is inversely correlated with the number of parts combined^33^, we reasoned that simplifying the final library assembly step to a two-fragment assembly is likely to result in higher quality final libraries. In addition, to facilitate introduction of the YFP dropout part in the second step, we took advantage of a previously-described methylation-switching approach^34^, which allows temporary blockage of SapI cut sites through the expression of a DNA methylase in *E. coli* (**Supplementary Figure 5A-B**).

To assess the composition of the ORI-marker libraries, we extracted plasmid DNA from the conjugation donors and performed amplicon sequencing (**Figure 3C** and **Supplementary Figure 6**). We found all ORIs were present in each of the six libraries, with 78.2% of all ORIs (108/138 ORIs across the six libraries) ranging in abundance between 0.5-10%. Certain ORIs were consistently overrepresented or underrepresented in all libraries: the pBBR1-UP, pNG2 and p15A ORIs together made up between 16.8-65.7% of library composition, whereas the pTHT15 ORI made up between 0.03-3.21% of libraries. It is important to note that quantification of library members from extracted plasmid DNA may not perfectly correlate to the abundance of cells harboring library variants, since some ORIs may increase or decrease the plasmid copy number within each conjugation donor cell (**Supplementary Figure 7**). Library members for which representation is reduced will be detected with lower sensitivity in ORI-marker screens, as the amount of DNA delivered will be decreased. However, we previously demonstrated detection of functional ORIs which made up as little as 0.11% of the total library^8^.

The POSSUM module provides all the components required to construct ORI-marker libraries in the manner described above. While we describe here the construction of only six libraries, each with a different antibiotic resistance gene, the module includes a total of 23 distinct antibiotic resistance markers allowing for a wide range of distinct libraries to be constructed for specific target microbes, including fungi (**Supplementary Table 6**). In addition, the modularity of the kit enables rapid generation of new ORI-marker combinations and allows users to customize the specific set of ORIs included in a given library for their particular use case, adding or removing ORIs as desired.

For users that do not wish to work with pooled plasmid libraries, ORI-marker variants can also be assembled individually; we include 17 such individually assembled ORI-marker constructs with a gentamicin resistance cassette and varying bacterial ORIs in the POSSUM module (**Supplementary Table 9**). Finally, to further facilitate library generation, the POSSUM module contains 11 pre-assembled library destination vectors (each with a different selection marker), 30 Level 1 barcoded ORI plasmids and the methylation-switchable YFP dropout part plasmid.

### MACKEREL module design and characterization

Central to many synthetic biology and genetic engineering approaches is the ability to drive the expression of recombinant proteins to desired levels. In the context of bacterial genetics, this typically means having a collection of transcriptional promoters and ribosome binding sites (RBSs) with well-characterized properties. However, the functional properties of such genetic tools can vary drastically from organism-to-organism, necessitating host-specific part characterization and screening^35,36^. To facilitate this process, we set out to create a broad host range promoter library that could be easily combined with the POSSUM module for use in non-model microbes.

We selected a well-characterized library of 421 promoter-RBS sequences from Yim et al.^23^, referred to as RS421. All 421 members of this library were derived from mobile genetic elements and demonstrate to be transcriptionally active to varying levels in cell-free lysates from 10 bacterial species spanning three phyla. We set out to generate an RS421 library Level 0 part, to enable Golden Gate assembly with POSSUM module components. First, internal BsaI and SapI sites were mutated and assembly overhangs were added to each RS421 variant. We also added four control promoter sequences spanning a range of strengths from the widely used Anderson promoter library^37^ (J23119, J23100, J23108 and J23114) and a non-functional dummy promoter. Since the promoter-RBS sequences are themselves short enough to be directly sequenced using Illumina sequencing platforms (165 bp), no unique barcodes were included. Finally, these 426 sequences were synthesized as an oligo pool and cloned into a Level 0 part storage vector creating the final pooled library which we dubbed the **MACKEREL module** (**Figure 4A**).

**Figure 4.**
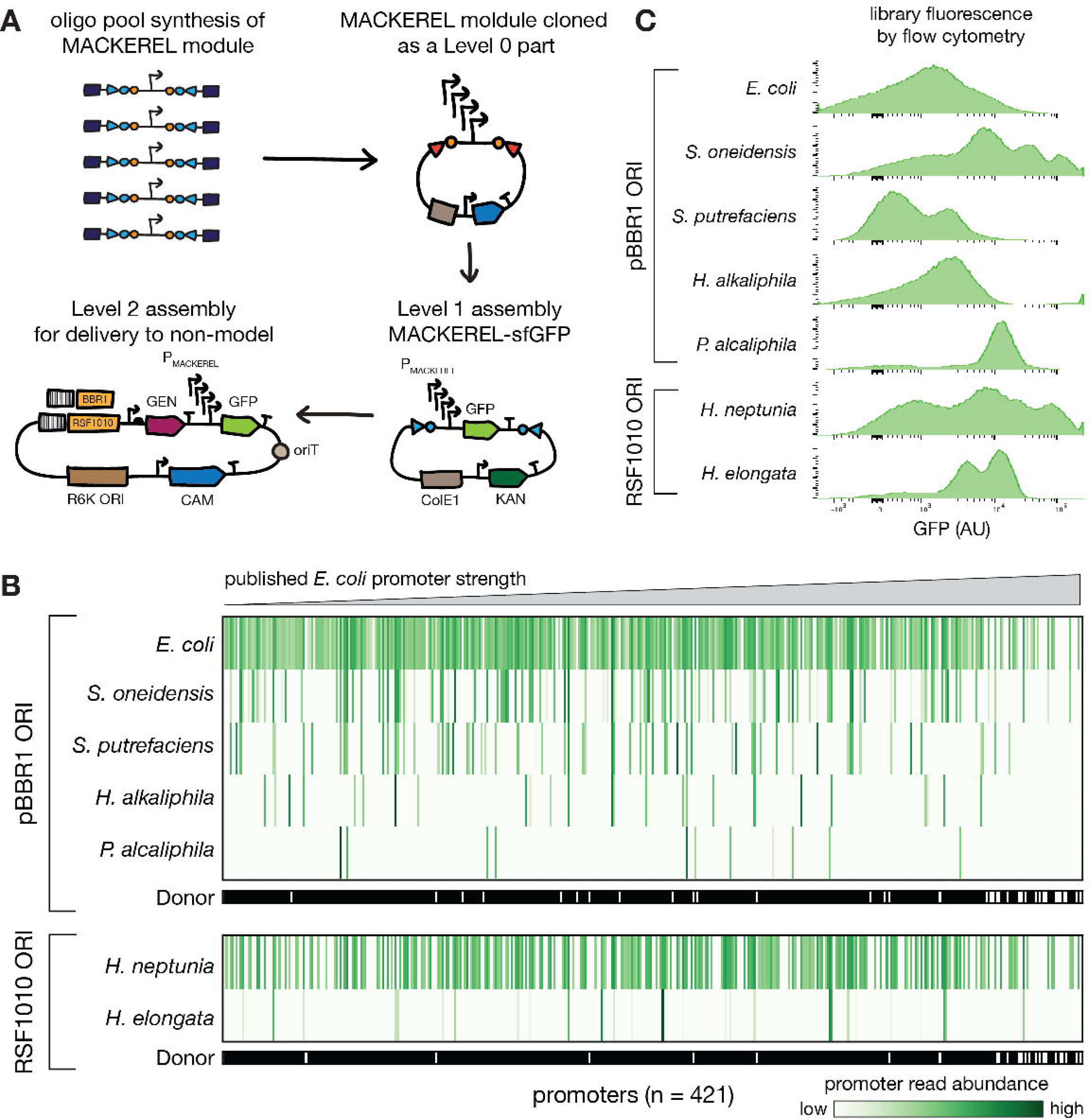
Assembly and screening of the MACKEREL module in non-model microbes. (A) Construction and application of the MACKEREL module. We synthesized the RS421 promoter library along with five control promoters as an oligo pool and cloned it into a Level 0 part storage vector, creating the MACKEREL module. Using the POSSUM module, we then combined the MACKEREL module with plasmid ORIs (pBBR1 and RSF1010), a gentamicin resistance marker (GEN), *oriT* and an sfGFP reporter gene to create libraries that could be delivered to diverse species. (B) Amplicon sequencing results showing normalized abundance of promoter reads in seven recipient organisms. The upper five recipient organisms were conjugated with the MACKEREL module-sfGFP cassette hosted on the pBBR1 backbone, while the lower two recipient organisms were conjugated with the MACKEREL module-sfGFP cassette hosted on the RSF1010 backbone. Promoters on the x-axis are sorted by their reported transcription strength for *E. coli* by Yim et al. (2019)^23^, excluding the 5 control promoters. The presence/absence of promoters in respective *E. coli* donor strain libraries is indicated in black/white, respectively, below. (C) Normalized fluorescence intensity (GFP) of pooled recipient transconjugant libraries as measured by flow cytometry. Flow cytometry was used to determine the distribution of fluorescence intensity across the entire library with the FITC fluorochrome, representing the range of GFP expression strengths driven by MACKEREL module variants. *E. coli* (*Escherichia coli* Zymo 10B), *S. oneidensis* (*Shewanella oneidensis* MR-1), *S. putrefaciens* 95 (*Shewanella putrefaciens* 95), *H. alkaliphila* (*Halomonas alkaliphila* 18b Ag), *P. alcaliphila* (*Pseudomonas alcaliphila* AL 15-21), *H. neptunia* (*Halomonas neptunia* A1), *H. elongata* (*Halomonas elongata* 1H9).

To determine the abundance of promoter variants in the MACKEREL module, we performed amplicon-sequencing in the *E. coli* donor strain and corroborated this with whole plasmid sequencing, confirming that all 426 variants were present in the library, albeit with variable abundance (**Supplementary Figure 8**). Notably, a small number of promoter library variants had significantly lower or zero abundance in the amplicon-sequencing dataset compared to the whole plasmid sequencing dataset. In these cases, we suspect sequence-specific effects may have resulted in inefficient PCR-amplification of certain variants. Promoter abundance was inversely correlated with published *E. coli* transcription strength scores^23^, suggesting that high strength promoters impose a metabolic burden to host cells (**Supplementary Figure 9**).

### Application of the MACKEREL module gene expression in non-model microbes

We next set out to screen MACKEREL module variants for their ability to drive gene expression in a panel of bacteria from the phylum Pseudomonadota. To achieve this, we used Golden Gate assembly to combine the MACKEREL module with components from the POSSUM module to allow transformation to and characterization in non-model bacteria.

First, the MACKEREL module was assembled with a fluorescent reporter CDS (sfGFP) and transcriptional terminator (**Figure 4A** and **Supplementary Figure 10**). This reporter gene cassette was then assembled with either of two broad host range plasmid ORIs (pBBR1 and RSF1010), a gentamicin resistance cassette and conjugation *oriT* sequence (**Figure 4A** and **Supplementary Figure 11**). Notably, we had previously determined the gentamicin resistance cassette and plasmid ORIs used here were functional in these species using our ORI-marker screen approach^8^. Since Golden Gate assembly transformations usually contain a non-negligible proportion of re-ligated destination vector, we created a set of 20 destination vectors in which the dropout part contained a *ccdB* cassette (**Supplementary Table 2**). Plasmids encoding the *ccdB* toxin can only be maintained in specific cloning strains, such as ccdB Survival 2 T1R *E. coli* (Invitrogen). Consequently, library assemblies were transformed into *ccdB* non-permissive *E. coli*, preventing maintenance of re-ligated destination vectors. Finally, libraries were transformed into an *E. coli* conjugation donor strain and delivered to seven target bacterial species from the phylum Pseudomonadota: *E. coli*, *Shewanella oneidensis*, *Shewanella putrefaciens*, *Pseudomonas alcaliphila*, *Halomonas alkaliphila*, *Halomonas neptunia* and *Halomonas elongata*.

To determine library composition in both the *E. coli* conjugation donor and recipient species transconjugant libraries, we performed targeted amplicon-sequencing of the sfGFP promoter region (**Figure 4B**). We found both *E. coli* donor libraries possessed >90% of promoter variants (pBBR1 ORI library contained 385 of 426 promoter variants (90.3%); RSF1010 ORI library contained 397 of 426 promoter variants (93.1%)). The high diversity of promoter variants found in the donor underscores the robustness of our library assembly and delivery protocols. The number of transconjugant colony-forming units (CFU) obtained varied over several orders of magnitude between recipient species, likely due to variation in the conjugation efficiency, from a maximum of 2.1 x 10^7^ CFU for *E. coli* to 20 CFU for *H. alkaliphila* (**Supplementary Figure 12A**). As expected, we found the number of promoters detected by NGS in each recipient transconjugant library was positively correlated with the number of transconjugant CFU obtained (**Supplementary Figure 12B**), highlighting the importance of efficient DNA delivery methods for library screening.

Next, to assess the range of expression strengths driven by MACKEREL module variants, transconjugant libraries were analyzed by flow cytometry (**Figure 4C**) and fluorescence imaging of colonies on solid media (**Supplementary Figure 13**). Both methods revealed a wide range of GFP-expression intensities among transconjugants. These results are consistent with data reported by Yim et al.^23^, which illustrate that RS421 library variants are able to direct gene expression over a range of strengths across at least three bacterial phyla. Combining the MACKEREL module with POSSUM module components therefore enables rapid, simple screening of gene expression elements in non-model bacteria.

Although beyond the scope of this study, approaches such as FACS-Seq could be applied to transconjugant libraries to determine and compare the strength of protein expression driven by individual MACKEREL module members for different non-model microbes^2^. Further, the modularity of Golden Gate assembly allows users to deploy the MACKEREL module Level 0 part for a variety of different applications, for instance in functional screens for optimal expression of a recombinant protein or of metabolic pathways. To support these and other applications, the MACKEREL module Level 0 part promoter library is being made available as a pooled library through Addgene.

## Conclusion

Here we have described two resources designed to support the identification of foundational genetic tools in non-model microbes. The POSSUM module enables modular assembly of genetic parts and identification of functional ORIs and selection markers. The MACKEREL module enables identification of functional expression elements across a large dynamic range in non-model microbes. By combining the modularity and flexibility of Golden Gate assembly with high-throughput screening methods such as NGS and flow cytometry, we provide a rapid and efficient way to identify functional genetic parts in under-studied microbes. For a given microbe, functional parts can then be reassembled with parts useful for specific applications such as CRISPR- or recombineering-based genome engineering, tuning secretion elements for recombinant protein production or expression of multi-gene metabolic pathways. The design framework presented here enables a scalable method for establishing key initial footholds in genetic tractability in non-model microbes, and is extensible for a growing library of genetic parts and for complex hierarchical assemblies. Further development and broad application of this pan-microbial toolkit will accelerate our ability to study and engineer diverse microbes.

## Materials and Methods

### Growth conditions

All strains used in this study were grown in LB broth or on LB 2% agar supplemented with diaminopimelic acid (DAP) 60 µg/mL, chloramphenicol (CAM) 20 µg/mL, carbenicillin (CARB) 100 µg/mL, kanamycin (KAN) 50 µg/mL, spectinomycin (SPEC) 100 µg/mL, gentamicin (GEN) 20 µg/mL and/or 2% (w/v) D-glucose when appropriate. Liquid cultures were grown with shaking at 225 RPM. The conjugation donor strain *Escherichia coli* BW29427 (*thrB1004 pro thi rpsL hsdS lacZΔM15 RP4-1360 Δ(araBAD)567 ΔdapA1341::[erm pir]*) was with DAP supplementation at all times. Details of all bacterial strains and their respective growth temperatures are summarized in **Supplementary Table 10**.

### Design and construction of POSSUM module Golden Gate assembly destination vectors

Golden Gate assembly destination vectors created in this study follow the architecture of Loop Assembly destination vectors^15,25^. All destination vectors shared the following features: an *E. coli* ORI (ColE1 or R6K); an *E. coli* antibiotic resistance cassette (CAM, KAN or AMP); internal SapI/BsaI cut sites for Golden Gate assemblies into the vector; external BsaI/SapI cut sites for assemblies out of the vector; unique nucleotide sequences (UNSs) flanking the assembly cargo that can also be used as long overlaps for the cloning of transcription units into larger constructs^25,38^; transcription terminators (L2U5H11 and L2U8H11^39^) flanking the assembly site to prevent unintentional cargo expression; a fluorescent protein (sfGFP or mScarlet-I) dropout part which is replaced by the assembly cargo allowing easy discrimination of re-ligated destination vector on transformation plates^40^. Notably, the fluorescent protein expression cassettes used within dropout parts employ a very strong *E. coli* promoter (PglpT). To reduce the risk of mutations arising in this cassette, when strains hosting these plasmids were grown routinely, culture medium was supplemented with 2% (w/v) glucose which represses expression from the PglpT promoter^41^.

All Golden Gate assembly destination vectors possessing the ColE1 ORI (pGLDV_1-17, pGLDV_21) were generated by Clonal Gene Synthesis (Twist Biosciences). The standard pTwist High Copy cloning vectors provided the ColE1 ORI and AMP/KAN/CAM markers while custom-synthesized regions contained the SapI/BsaI sites, UNSs, terminators and dropout parts. For plasmids pGLDV_19, 20, 41 and 42, the ColE1 ORI was replaced with the R6Kγ ORI using NEBuilder HiFi DNA Assembly (New England Biolabs). The R6Kγ ORI was amplified from pBAM1^42^, NEB HiFi assemblies were performed according to the manufacturer’s instructions and assembly products were transformed into TransforMax™ EC100D™ pir-116 *E. coli* (Biosearch Technologies).

Plasmids pGLDV_43-58 were created by replacing the sfGFP dropout part in the corresponding plasmids pGLDV_1-16 with an mScarlet-I-*ccdB* dropout part synthesized by Custom Gene Synthesis (Twist Biosciences) in BsaI or SapI Golden Gate assembly reactions. Assembly products were transformed into ccdB Survival 2 T1R *E. coli* (Invitrogen). Plasmids pGLDV_59-63 were created by replacing the sfGFP dropout part in the corresponding plasmids pGLDV_19, 20, 41 and 42 with an ColE1-mScarlet-I-*ccdB* dropout part synthesized by Custom Gene Synthesis (Twist Biosciences) in BsaI or SapI Golden Gate assembly reactions. Assembly products were transformed into ccdB Survival 2 T1R *E. coli* (Invitrogen). Since no *ccdB*-resistant, pir-plus strains were commercially available, we introduced a ColE1 ORI into the dropout part of these four plasmids.

### Construction of POSSUM module Level 0 part plasmids

Details of the construction method used for all Level 0 plasmids are listed in **Supplementary Table 4**. Where possible, Level 0 part sequences were commercially synthesized as sequence-verified clonal genes into high copy backbones, with standard Level 0 vector features added (BsaI sites and overhangs, flanking terminators, UNSs). However, in many cases commercial gene synthesis was not possible as sequence complexity filters would not accept repetitive and high/low GC-content fragments, which are particularly common in plasmid ORIs. These parts were constructed by obtaining a template plasmid from Addgene, amplifying the desired fragment by PCR with primers that introduce the required SapI cut sites and overhangs, and assembling into the pGLDV_17 destination vector. In cases where parts contained internal SapI or BsaI sites that needed to be mutated, sequences were amplified in multiple fragments with fragment boundaries at the mutation sites. When SapI/BsaI sites were within coding sequences, primers were designed to create silent mutations. When SapI/BsaI sites were within intergenic regions, primers were designed to create mutations that preserved GC-content. All Level 0 part plasmids were confirmed by Sanger sequencing of the part region (Eton Bioscience, Inc.) or by whole plasmid sequencing (Primordium Labs). In addition, all plasmids deposited to Addgene were independently subjected to whole-plasmid sequence confirmation by Addgene.

### Design and construction of miscellaneous POSSUM module plasmids

To cater for instances where fewer than four parts must be combined in Odd/Even assemblies, we created a set of six end-linker plasmids (**Supplementary Table 11**). These end-linker parts consist of short (40 bp), non-functional sequences flanked by BsaI/SapI cut sites that allow them to occupy vacant assembly positions. To create end-linker parts we used six biologically inactive UNSs from Torella et al. (2014)^38^. Clonal Gene Synthesis (Twist Bioscience) was used to synthesize end-linker parts with the appropriate flanking SapI/BsaI cut sites and plasmid backbones.

The methylation-switchable sfYFP dropout part plasmid (pGLDO_1) was generated by Clonal Gene Synthesis (Twist Bioscience) (**Supplementary Table 11**). The sfYFP cassette possessed identical promoter and terminator sequences to the sfGFP dropout part cassette described above. Methylation-switchable SapI sites were designed following the MetClo system^34^. For methylation-based blockage of SapI sites, the pGLDO_1 plasmid was transformed into *E. coli* DH10B M.XmnI (Addgene, 114269)^34^ and plasmid DNA was purified using the QIAprep Spin Miniprep Kit (Qiagen, 27104).

### Standard Golden Gate assemblies

Standard Golden Gate assemblies were performed using a protocol adapted from Pollak et al. (2019)^15^ with total reaction volumes scaled down from 10 µL to 5 µL. For all assemblies, a 2.5 µL DNA mix was first prepared in PCR strip tubes. This mix contained 0.5 µL of each part and destination vector plasmid, which were previously diluted to 50 nM concentration, and the final volume was made up to 2.5 µL with H_2_O. Next SapI/BsaI enzyme master mixes were prepared on ice (**Supplementary Table 12**), 2.5 µL master mix was combined with each DNA mix and aspirated to thoroughly combine. Reactions were then incubated using the thermocycler programs described in **Supplementary Table 13**. Where possible, assembly products were transformed into *E. coli* Mix & Go! Competent Cells-Zymo 10B (Zymo Research, T3020). For assemblies using destination vectors possessing the R6Kγ ORI, products were transformed into chemically-competent TransforMax™ EC100D™ pir-116 *E. coli* (Biosearch Technologies), prepared using the Mix & Go! *E. coli* Transformation Kit and Buffer Set (Zymo Research, T3002). For assemblies requiring maintenance of the *ccdB* cassette, products were transformed into One Shot™ ccdB Survival™ 2 T1R Competent Cells (Invitrogen, A10460).

### Pooled ORI-marker library Golden Gate assemblies

To generate pooled ORI-marker libraries, Level 1 barcoded-ORI plasmids were first manually pooled by combining equal volumes of 50 nM plasmid stocks, with the exception of the dummy ORI plasmid (pGL1_46) for which double the volume was used. Since the fold enrichment in NGS reads mapping to each ORI compared to the dummy ORI is used to assign functional ORIs in the ORI-marker screen^8^, maintaining the dummy ORI at a relatively high abundance in ORI-marker libraries helps reduce the risk of false positives. Next, SapI Golden Gate assemblies were performed as described in the previous section treating the pooled barcoded-ORI as a typical assembly part. To enable electroporation, completed Golden Gate assembly reactions were purified using the DNA Clean & Concentrator-5 (Zymo Research, D4013) with final elution in 6 µL molecular biology grade H_2_O. Next, 2 µL of purified Golden Gate assembly mix was transformed into *E. coli* BW29427 electrocompetent cells. After 1 hour recovery at 37°C, 100 µL of 10^-3^ and 10^-5^ dilutions were plated onto selective media for determination of total number of transformants. The remaining 950 µL of recovered cells were inoculated into 100 mL of liquid selective media and grown for 16 hours at 30°C. Plasmid DNA was extracted from 5 mL of this culture using the QIAprep Spin Miniprep Kit (Qiagen, 27104). The remaining 95 mL of culture was combined with 40.7 mL 50% glycerol (v/v), aliquoted into 1 mL stocks and stored at −80°C for use in ORI-marker screens.

### Design and construction of the MACKEREL module

The sequences of the RS421 promoter-RBS library were obtained from Yim et al. (2019)^23^. Five internal control promoters with known relative expression strengths in *E. coli* were also selected - a 48-bp randomly-generated non-functional dummy promoter sequence and four Anderson library promoters^37^: J23114 (low strength), J23108 (medium strength), J23100 (high strength) and J23119 (very high strength). A strong bacterial ribosome binding site (BBa B0034^43^) was added to each of the control promoters and overall length was adjusted to 165 bp to match the length of the RS421 variants. To make parts compatible with Golden Gate assembly, internal BsaI/SapI sites were mutated to remove cut sites but preserve GC-content and upstream and downstream SapI sites and overhangs were added. Finally, primer binding sites used by Uzonyi et al. 2022^44^ were added to enable PCR amplification of the entire pooled library following commercial synthesis. Oligos were ordered as a pooled library from Twist Biosciences. Upon receipt, oligos were amplified according to Uzonyi et al. 2022^44^ and assembled by Golden Gate (SapI) into the Level 0 part storage vector pGLDV_17, as described below.

### MACKEREL module Golden Gate assemblies

To generate the MACKEREL module Level 0 part (pGL0_151), the PCR-amplified oligo pool was first assembled into the pGLDV_17 Level 0 part storage plasmid by standard SapI Golden Gate assembly scaled up to a 20 µL final volume. To enable electroporation, completed Golden Gate assembly reactions were purified using the DNA Clean & Concentrator-5 (Zymo Research, D4013) with final elution in 10 µL molecular biology grade H_2_O. Next, 1 µL of purified Golden Gate assembly mix was transformed into NEB 10-beta Electrocompetent *E. coli* (New England Biolabs, C3020K) according to the manufacturer’s instructions. After a 1 hour recovery, 100 µL of 10^-2^ and 10^-4^ dilutions were plated onto selective media for determination of total transformants, which showed a per-variant coverage of >10^4^-fold. The remaining 950 µL of recovered cells were inoculated into 50 mL of liquid selective media and grown for 6.5 hours at 37°C shaking at 225 rpm (OD600 = 1.85). Plasmid DNA was extracted from ten 4-mL aliquots of this culture using the QIAprep Spin Miniprep Kit (Qiagen, 27104).

To create a MACKEREL module-sfGFP Level 1 cassette (pGL1_75), pGL0_151 was assembled with the required Level 0 parts (**Supplementary Table 8**) by standard BsaI Golden Gate assembly scaled up to a 20 µL final volume. To enable electroporation, completed Golden Gate assembly reactions were purified using the DNA Clean & Concentrator-5 (Zymo Research, D4013) with final elution in 6 µL molecular biology grade H_2_O. Next, 2 µL of purified Golden Gate assembly mix was transformed into NEB 10-beta Electrocompetent *E. coli* (New England Biolabs, C3020K) according to the manufacturer’s instructions. After a 1 hour recovery, 100 µL of 10^-2^, 10^-4^ and 10^-6^ dilutions were plated onto selective media for determination of total transformants, which showed a per-variant coverage of >10^3^-fold. The remaining 950 µL of recovered cells were inoculated into 50 mL of liquid selective media and grown for 6.5 hours at 37°C shaking at 225 rpm (OD600 = 1.06). Plasmid DNA was extracted from five 10-mL aliquots of this culture using the QIAprep Spin Miniprep Kit (Qiagen, 27104).

To enable conjugation of the sfGFP cassette to non-model bacteria, pGL1_75 was combined with the RSF1010 ORI or pBBR1 ORI, a gentamicin resistance cassette and the RP4/RK2 *oriT* sequence by standard SapI Golden Gate assembly, creating pGL2_202 (RSF1010) and pGL2_204 (pBBR1). To enable electroporation, completed Golden Gate assembly reactions were purified using the DNA Clean & Concentrator-5 (Zymo Research, D4013) with final elution in 6 µL molecular biology grade H_2_O. Next, 1 µL of purified Golden Gate assembly mix was transformed into NEB 10-beta Electrocompetent *E. coli* (New England Biolabs, C3020K) according to the manufacturer’s instructions. After a 1 hour recovery, 100 µL of 10^-2^ and 10^-4^ dilutions were plated onto selective media for determination of total transformants, which showed a per-variant coverage of >10^3^-fold. The remaining 950 µL of recovered cells were inoculated into 50 mL of liquid selective media and grown for 7.5 hours at 37°C shaking at 225 rpm. Plasmid DNA was extracted from this culture using the QIAprep Spin Miniprep Kit (Qiagen, 27104). Finally, extracted plasmid DNA was transformed into *E. coli* BW29427 electrocompetent cells. After a 1 hour recovery, 100 µL of 10^-1^ and 10^-2^ dilutions were plated onto selective media for determination of total transformants. The remaining 950 µL of recovered cells were inoculated into 50 mL of liquid selective media and grown for 7.5 hours at 37°C. Plasmid DNA was extracted from 5 mL of this culture using the QIAprep Spin Miniprep Kit (Qiagen, 27104). Glycerol was added to the remaining culture to a final concentration of 15% (v/v) and the resulting mix stored at −80°C.

For both Level 1 (pGL1_75) and Level 2 (pGL2_202, 204) assemblies, library transformants on solid media were imaged for fluorescence using GFP imaging conditions on a Phenobooth+ (Singer Instruments) system (excitation LED: λ = 390 nm peak, 375-400 nm 50% intensity range; emission filter λ = 527 nm).

### Sequencing-based quantification of libraries

Amplicon sequencing library preparation was performed as previously described (details available at Gilbert et al.^8^), using primers designed to amplify the barcodes associated with both ORI and promoter sequences. Briefly, extracted plasmid libraries were diluted to 0.5 ng/µL in Buffer EB (Qiagen, 19086) and Kapa Hifi HotStart Ready mastermix (Roche) was used for the PCR reactions with the following primer sets (**Supplementary Table 14**). The concentration of the resultant amplicons were measured using a Qubit™ Flex Fluorometer (Thermo Fisher Scientific) and each sample was diluted to 1 ng/µL for ORI-marker sequencing or by 50X for promoter library sequencing. Barcoded samples were pooled separately for ORI-marker and promoter library samples, but with equal volume, then purified with DNA Clean & Concentrator-5 (Zymo Research). The concentration of the purified, barcoded sample was measured using Qubit™ Flex Fluorometer (Thermo Fisher Scientific), and the DNA fragment size of the amplicon was confirmed to have the expected size of 216 bp (ORI-marker) or 384 bp (promoter library) using Invitrogen E-Gel EX agarose gel electrophoresis. The rest of the library preparation was completed following the Denature and Dilute Libraries Guide for MiniSeq System by Illumina, then loaded and sequenced on Illumina MiniSeq with a 300-cycle kit (2 x 150 bp paired-end run).

To estimate the abundances of either promoters or plasmid ORIs in a given sample, we quantified the number of reads matching the respective barcode associated with each part as previously described^8^. Briefly, amplicon sequencing reads (paired 2 × 150 bp) were quality and adapter trimmed using BBDuk, and then merged with BBmerge. Trimmed and merged reads were matched against a database of barcode reference sequences associated with each ORI and promoter using vsearch30 with the setting search_exact, retaining only perfect barcode matches.

We further estimated the abundances of promoters in input libraries (i.e. those hosted in the *E. coli* BW29427 conjugation donor strain) using whole plasmid sequencing, and saw good correspondence with amplicon results (**Supplementary Figure 8**). Whole plasmid pool DNA was sequenced on an Illumina MiniSeq with a 300-cycle kit (2 x 150 bp paired-end), and sequencing reads were quality and adapter trimmed using BBDuk. Reads were mapped to a combined index of all plasmid variants in the pool. A custom Python script using the Pysam package v0.2 was used to count only reads with a mapping quality score > 2, mapped with at least 50 bp and 1 or fewer mismatches that mapped to only the unique region of each variant in the plasmid pool. Mean depth of coverage of the unique promoter region was then used to estimate sequence abundances.

### Conjugation of MACKEREL-sfGFP libraries to non-model microbes

Conjugative donor libraries *E. coli* BW29427 pGL2_202 and pGL2_204 were grown-up by inoculating 1 mL of frozen library stock into 50 mL of LB media supplemented with DAP (60 µg/mL) and CAM (34 µg/mL) and incubating for 2.25 h at 37°C shaking at 225 rpm. Recipient strains were either grown in LB media or on LB agar (**Supplementary Table 10**). Recipients grown in liquid media were grown overnight in 50 mL of LB media at 30°C/37°C shaking at 225 rpm to an OD600 between 0.120-0.584. The cultures were centrifuged at 4,000 x *g* at 4°C for 10 min. The supernatant was discarded, cell pellets were resuspended in 1 mL 1X PBS, transferred to 1.5 mL tubes and centrifuged at 12,000 rpm for 5 min at room temperature. The supernatant was again discarded and pellets were resuspended to a final OD600 ∼100 in 1X PBS. Recipients grown on LB agar were scraped directly from the fresh streak plate, inoculated into 1 mL of 1X PBS and adjusted to a final OD600 ∼100 in 1X PBS. To initiate conjugation, 50 µL of donor and 50 µL of recipient were mixed directly on an LB agar plate supplemented with DAP 60 µg/mL using a spreader until dry. To improve cell-to-cell contact between donor and recipient, conjugation plates were dried aseptically for 1 hour prior to conjugation. Conjugation was allowed to proceed by incubating plates at 30°C for 3 h.

Following conjugation, cells were harvested by scraping with 1 mL of 1X PBS and transferred to a 1.5 mL tube. Cells were centrifuged at 12000 rpm for 5 min at room temperature, supernatant was discarded and pellets were resuspended with 1 mL of 1X PBS. This process was repeated once. Serial dilutions of resuspended cells from 10^0^ to 10^-5^ were prepared and 100 µL samples were plated onto selective LB agar plates with 20 µg/mL GEN and incubated at 30°C. In parallel, 500 µL of undiluted resuspended conjugation mix and 250 µL of 10^-1^ diluted conjugation mix were inoculated into 50 mL LB with 20 µg/mL GEN liquid cultures and incubated at 30°C with shaking at 225 rpm. Once colonies had formed on agar plates, colony counts were obtained to determine total library size and plates were imaged for fluorescence using GFP imaging conditions on a Phenobooth+ (Singer Instruments) system (excitation LED: λ = 390 nm peak, 375-400 nm 50% intensity range; emission filter λ = 527 nm). All plate images shown were generated under identical imaging settings. Once turbidity was observed in liquid cultures, samples were back-diluted 100-fold in fresh LB GEN 20 µg/mL liquid media and grown for ∼2 h prior to FACS analysis as described below. Plasmid DNA extractions were performed on cell pellets from 15 mL of these back-diluted cultures using QIAprep Spin Miniprep Kit (Qiagen, 27104).

### Measurement of MACKEREL-sfGFP library fluorescence by FACS and imaging

Fluorescence distributions of the unsorted transconjugant libraries were visualized by flow cytometry through the FITC fluorochrome after dilution at room temperature in LB with GEN as described above. All flow cytometry data acquisition and sorting were carried out on a BD FACS Aria II. The distributions were analyzed with FlowJo v10.9. All the points present in the distributions were first gated for cells on the FSC/SSC (which defines size and intracellular concentrations).

### Preparation of electrocompetent *E. coli*

A single colony of *E. coli* BW29427 was inoculated into 50 mL of LB media supplemented with DAP 60 µg/mL and grown overnight at 37°C at 225 rpm. The next day, 5 mL of the saturated overnight culture was diluted into two 2-liter flasks containing 500 mL of LB media to reach a starting OD600 of 0.05. The cultures were grown for 2.5 hours to reach a final OD600 between 0.45-0.5. The flasks were incubated on ice for 30 minutes with occasional swirling to ensure even cooling. The cultures were transferred to ice-cold 500 mL centrifuge bottles and harvested by centrifugation at 4000 x *g* for 10 min at 4°C. The cell pellets were each washed by gentle swirling with 1) 500 mL of ice-cold pure H_2_O, 2) 250 mL of ice-cold pure H_2_O, and 3) 10 mL of ice-cold 10% glycerol. Between each wash, cells were harvested by centrifugation as before and the supernatant was decanted. Following the final wash, the cells were harvested by centrifugation, the supernatant was decanted and cells were resuspended in 1 mL of ice-cold 10% glycerol. Finally, 100 μL aliquots were transferred to 1.5 mL tubes, flash frozen in dry ice and stored at −80°C.

## Supporting information

Plasmid sequences

## Data and Resource Availability

POSSUM module components are available via Addgene. A set of 189 plasmids are available to order as a single kit (<u>POSSUM Toolkit</u>, Kit ID #1000000234), while additional plasmids can be ordered individually (POSSUM Toolkit Expansion Pack), see **Supplementary Table 15** for further details. Annotated plasmid sequences for all POSSUM module plasmids are available as GenBank files on the bioRχiv webpage. Sequencing analysis scripts and data for POSSUM and MACKEREL screens are available at https://github.com/cultivarium/ORI-marker-screen/.

## Author Contributions

C.G., N.O. and H.H.L. conceived of and designed the project. C.G. performed plasmid design. C.G. and C.K. performed plasmid cloning. S.B. performed bacterial conjugations. E.L.D. performed flow cytometry experiments. C.K. performed DNA sequencing workflows. A.C.C. analyzed sequencing data.

C.G., S.B., N.O., A.C.C., E.L.D., C.K. and H.H.L wrote the manuscript. All authors have read and approved the manuscript.

## Acknowledgements

We thank all present and former members of the Cultivarium team for helpful discussions, technical advice and support throughout this project as well as helpful comments on this manuscript: Allison Lord, James Knight, Michael Molla, Mary-Anne Nguyen, Kerrin Mendler, Wajd Alsharif, Ariela Esmurria, Zaira Martín Moldes, Tyler Barnum, Julia Leung, Lilan Ling, Brianna Connolly, Julian Barros, Anna Douyon and Sierra Harken. Cultivarium acknowledges support from Schmidt Futures as a Convergent Research Focused Research Organization (FRO).

