## Supplementary Figures for "Design and construction towards a pan-microbial toolkit"

**Supplementary Figures** **2**

Supplementary Figure 1. Schematic comparing the architecture of the uLoop Assembly system to other toolkits. 2

Supplementary Figure 2. Assembly destination vectors included in the POSSUM module. 3

Supplementary Figure 3. Assembly compatibility of the full set of Level 0 parts in the POSSUM module. 5

Supplementary Figure 4. Level 1 assemblies performed to generate ORI-marker libraries. 6

Supplementary Figure 5. Schematic illustrating construction of pre-assembled ORI-marker destination vectors. 7

Supplementary Figure 6. ORI-marker library composition. 9

Supplementary Figure 7. Schematic illustrating the effect of plasmid copy number on library quantification through a hypothetical example. 10

Supplementary Figure 8. MACKEREL module Level 0 part composition. 11

Supplementary Figure 9. Correlation between MACKEREL module promoter abundance and transcription strength in E. coli. 12

Supplementary Figure 10. Level 1 assembly of the MACKEREL module with sfGFP. 13

Supplementary Figure 11. Level 2 assemblies of the MACKEREL module-sfGFP cassette with components required for non-model microbe transformation. 14

Supplementary Figure 12. Characterization of MACKEREL module transconjugants. 15

(A) Table showing the number of transconjugant colony forming units (CFUs) obtained from delivery of MACKEREL module constructs to recipient species. (B) XY scatter plot comparing the number of promoter library transconjugants obtained for each recipient species against the number of promoter variants detected in transconjugant libraries by NGS. 15

Supplementary Figure 13. Fluorescence imaging of MACKEREL module transconjugant plates. 16

**Bibliography** **17**

### Supplementary Figures

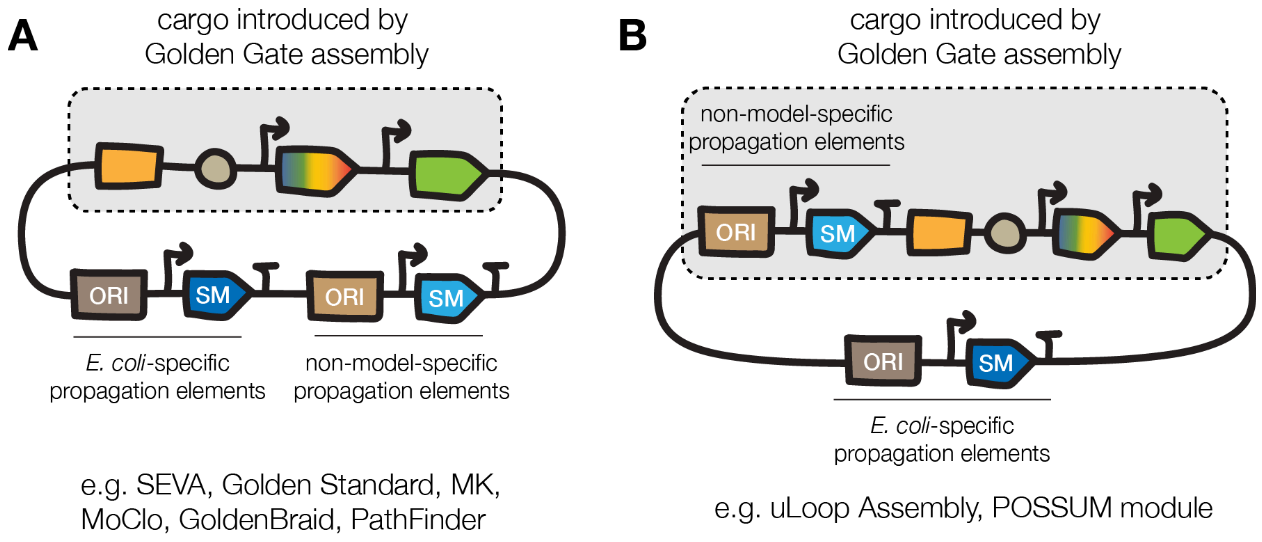


**Supplementary Figure 1. Schematic comparing the architecture of the uLoop Assembly system to other toolkits.**
(A) Most genetic toolkits designed for non-model microorganisms include known non-model-specific propagation elements on standard destination vector backbones. (B) The uLoop Assembly system includes non-model-specific propagation elements *within* the Golden Gate assembly cargo. This feature allows combinatorial assembly of ORIs and selection markers, which facilitates construction of ORI-marker test constructs. ORI: plasmid origin of replication sequence; SM: selection marker.

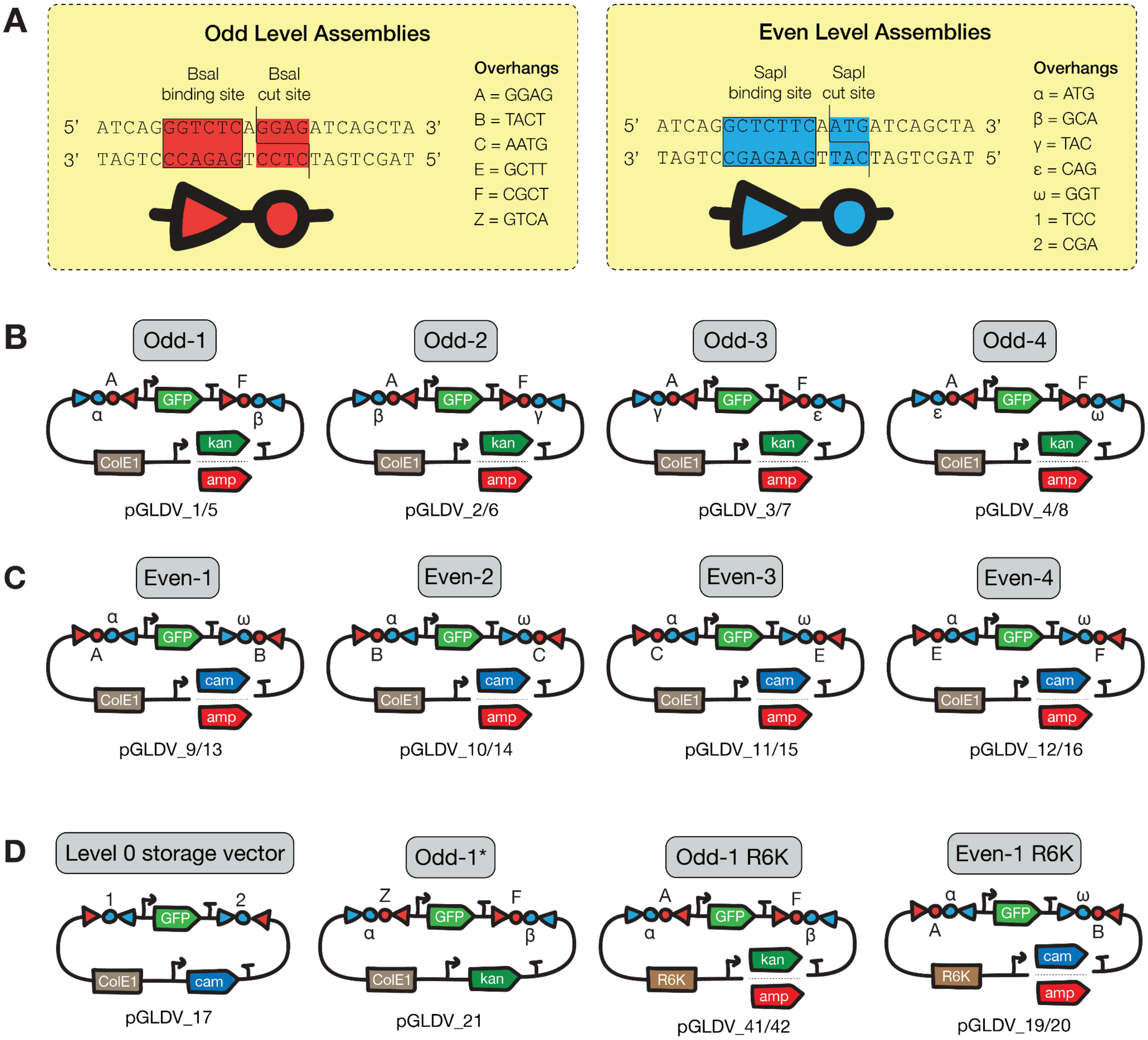


**Supplementary Figure 2. Assembly destination vectors included in the POSSUM module.** (A) Schematic of the "triangle-circle" notation used throughout this manuscript. Triangles represent Type IIS restriction enzyme binding sites and circles represent their cut sites. The location of the triangle apex represents the directionality of the enzyme binding site. BsaI sites are in red, SapI sites are in blue. Symbols A, B, C, E, F and Z represent unique four-base pair overhangs generated by BsaI. Symbols α, β, γ, ε, ω, 1 and 2 represent unique three-base pair overhangs generated by SapI (See Supplementary Figure 3). (B-C) Schematics of (B) Odd level and (C) Even level destination vectors included in the POSSUM module. We created two variants of each plasmid differing in the selection marker used. (D) Schematic showing non-standard destination vectors included in the POSSUM module. The Level 0 storage vector acts as an assembly destination vector for building new Level 0 parts. Odd-1* is an assembly destination vector which possesses an upstream Z overhang instead of A which therefore violates the common syntax. Odd-1 R6K and Even-1 R6K plasmids possess an R6Kγ origin of replication, which restricts their replication to engineered *E. coli* cloning strains that express the *pir* gene. We created two variants of the Odd-1 R6K and Even 1-R6K plasmids differing in the selection marker used. ColE1: high copy number *E. coli* ColE1 plasmid ORI; GFP: superfolder GFP under the very strong, glucose-repressible PglpT promoter[1]; kan: kanamycin resistance gene; cam: chloramphenicol resistance gene; amp: ampicillin/carbenicillin resistance gene; R6K: R6Kγ conditional replicon[2,3]. Plasmid names are indicated by pGLDV_X.

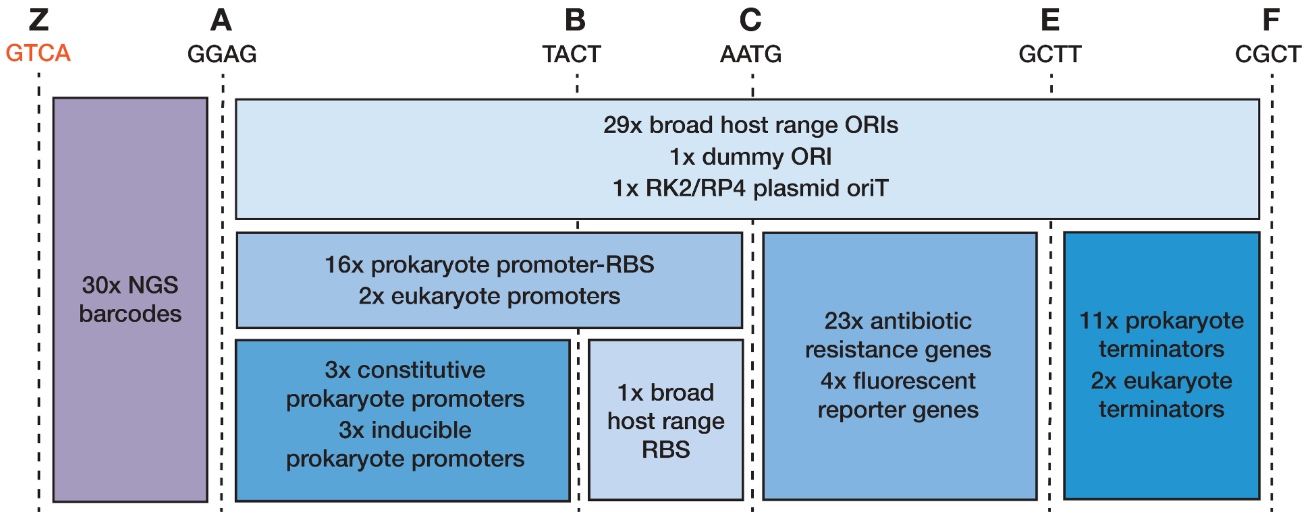


**Supplementary Figure 3. Assembly compatibility of the full set of Level 0 parts in the POSSUM module.**
This block view illustrates the assembly compatibility of Level 0 parts in the POSSUM module. Blocks represent assembly positions for subsequent Level 1 assemblies. Note that Level 1 assembly destination vectors are able to accept parts spanning overhangs A-to-F. Vertical dotted lines represent assembly boundaries, with the corresponding four base pair overhang and single letter code indicated at the top. The number and type of Level 0 parts filling each position is indicated within the colored-blocks. For instance, the POSSUM module contains 23x antibiotic resistance gene coding sequence Level 0 parts, each with an upstream AATG and downstream GCTT overhang. The Z (GTCA) overhang, highlighted in orange, can only be used in the non-standard assembly destination vector pGLDV_21.

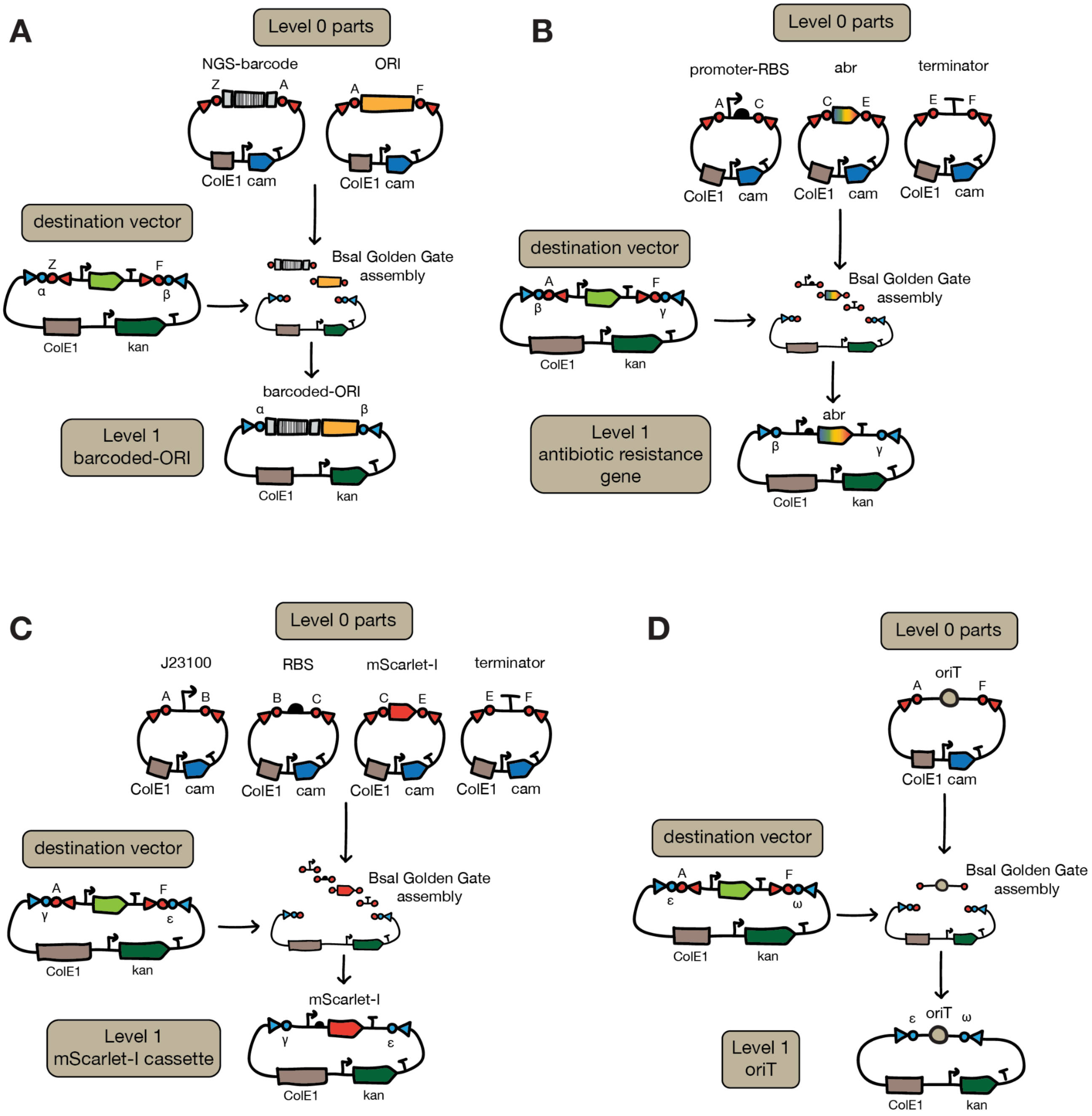


**Supplementary Figure 4. Level 1 assemblies performed to generate ORI-marker libraries.** (A) Level 0 plasmid ORI parts were combined with Level 0 NGS barcode parts and assembled into the non-standard Odd-1* destination vector (pGLDV_21). (B) Antibiotic resistance genes were re-created from their component Level 0 parts and assembled into an Odd-2 assembly destination vector. (C) An mScarlet-I expression cassette was assembled with a medium-to-strong constitutive promoter (J23100) into an Odd-3 assembly destination vector. (D) The *oriT* Level 0 part was assembled into an Odd-4 assembly destination vector.

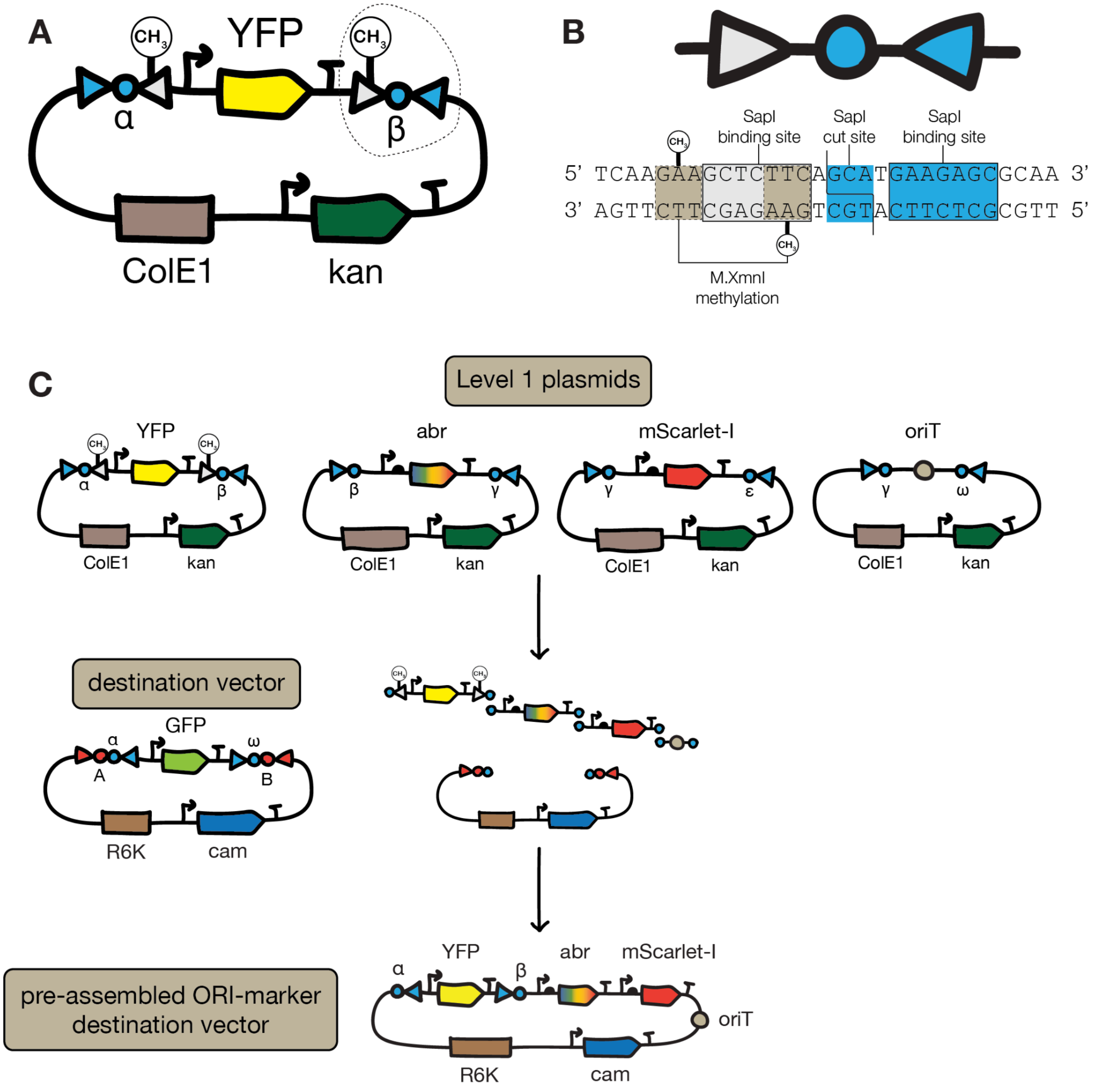


**Supplementary Figure 5. Schematic illustrating construction of pre-assembled ORI-marker destination vectors.**

(A) Generation of pre-assembled ORI-marker destination vectors relies on the use of a methylation-switchable YFP dropout part, which occupies the ORI position. Blue triangles represent standard SapI binding sites, blue circles represent the three base pair overhangs generated by SapI, gray triangles represent methylation-switchable SapI binding sites and $CH_3$ lollipops represent adenine methylation. (B) Detailed view of the sequence outlined in panel A, illustrating how methylation can block SapI cut sites in the YFP dropout part. The methylation-switchable SapI site (gray triangle) overlaps with the recognition sequence of M.XmnI methylase (GAANNNNTTC). When DNA is isolated from an *E. coli* strain engineered to express the

M.XmnI methylase, an adenine base within the SapI site is methylated. This methylation blocks SapI from binding and cutting. Note that the methylation-switchable (gray triangle) and standard (blue triangle) SapI sites are oriented so they both cut at the same position. (C) Level 1 plasmids were assembled into a Level 2 assembly destination vector possessing an R6K ORI. Notably, the YFP dropout part plasmid is extracted from an M.XmnI-expressing *E. coli* strain, meaning the methylation-switchable SapI sites are not cut during this assembly. The final assembly is transformed into an *E. coli* strain that lacks the M.XmnI methylase, meaning the methylation-switchable SapI sites can now be used for subsequent assemblies. This design therefore allows for creation of a new destination vector with pre-assembled antibiotic resistance gene, mScarlet-I gene and *oriT* sequence using a SapI Golden Gate assembly reaction as well as subsequent replacement of the YFP dropout part using another SapI Golden Gate assembly reaction (Main Figure 3B).

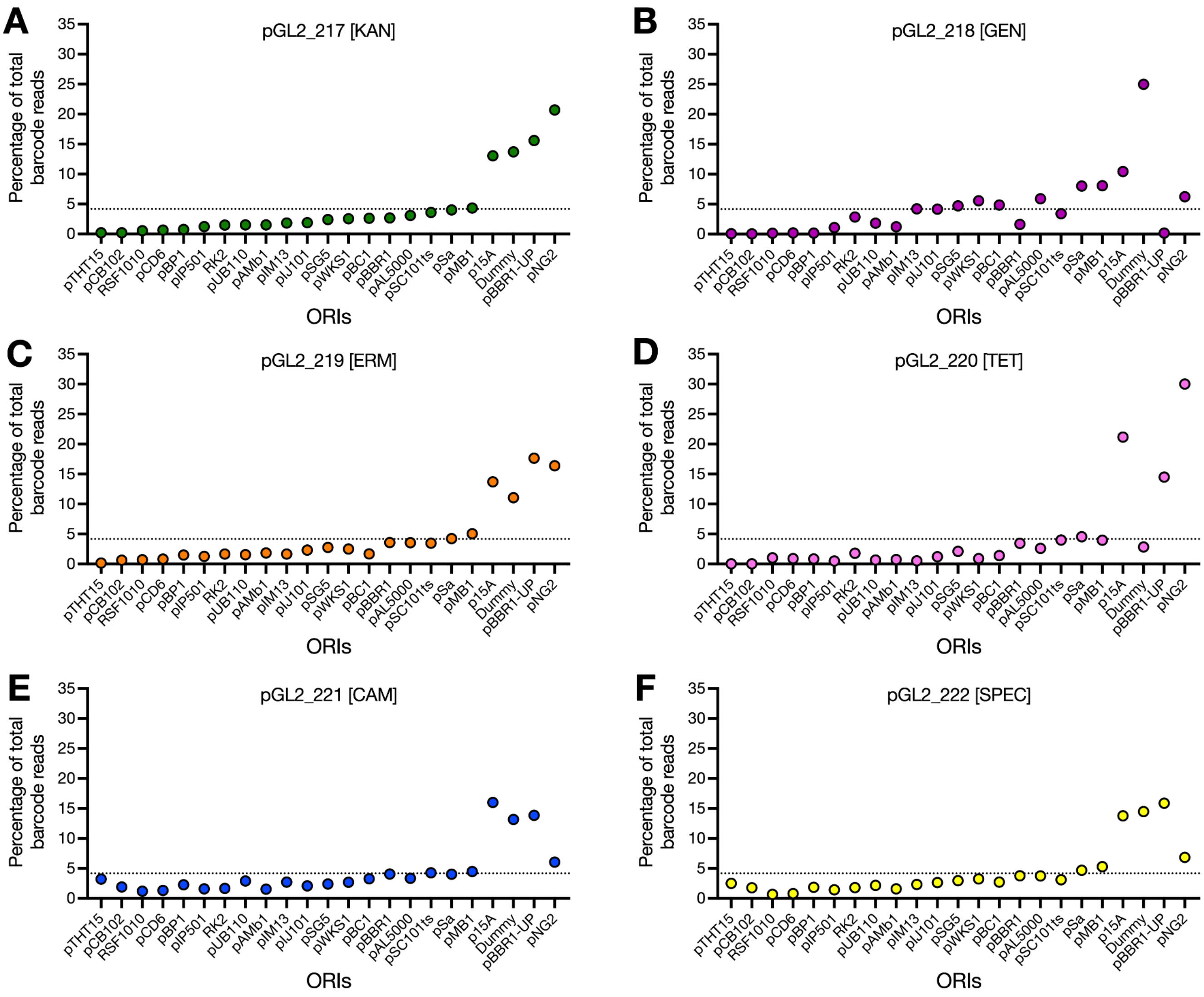


**Supplementary Figure 6. ORI-marker library composition.**
Scatter plots showing the percentage of total barcode reads mapping to each ORI barcode across six ORI-marker libraries extracted from conjugation donor strains: (A) pGL2_217 [KAN]; (B) pGL2_218 [GEN]; (C) pGL2_219 [ERM]; (D) pGL2_220 [TET]; (E) pGL2_221 [CAM]; (F) pGL2_222 [SPEC]. The horizontal dotted line represents the expected abundance of each ORI if library distribution was perfectly even.

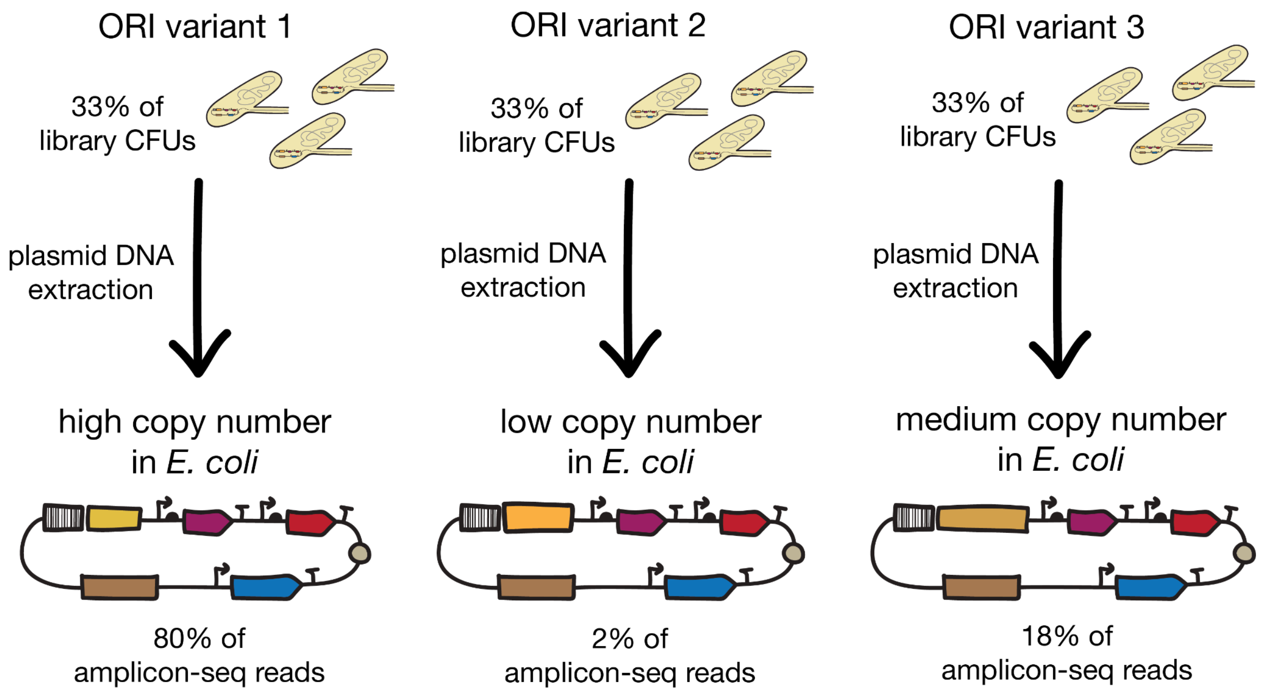


**Supplementary Figure 7. Schematic illustrating the effect of plasmid copy number on library quantification through a hypothetical example.**

In this hypothetical scenario, an ORI-marker library is composed of three different ORI variants. The transformed library has a perfectly even distribution, with colony forming units (CFUs) of each ORI variant making up one third (33%) of the total library population. Plasmid DNA is extracted and amplicon-sequencing performed. Since ORI variant 1 directed high copy number plasmid replication in *E. coli*, it is overrepresented in the extracted DNA library (80%) compared to the low copy number ORI variant 2 (2%) and medium copy number ORI variant 3 (18%). This is reflected in the abundance of reads generated by amplicon-sequencing. Notably, for DNA delivery by conjugation, the abundance of individual conjugation donor CFUs in the library is more important than the abundance of plasmid DNA per cell (i.e. copy number).

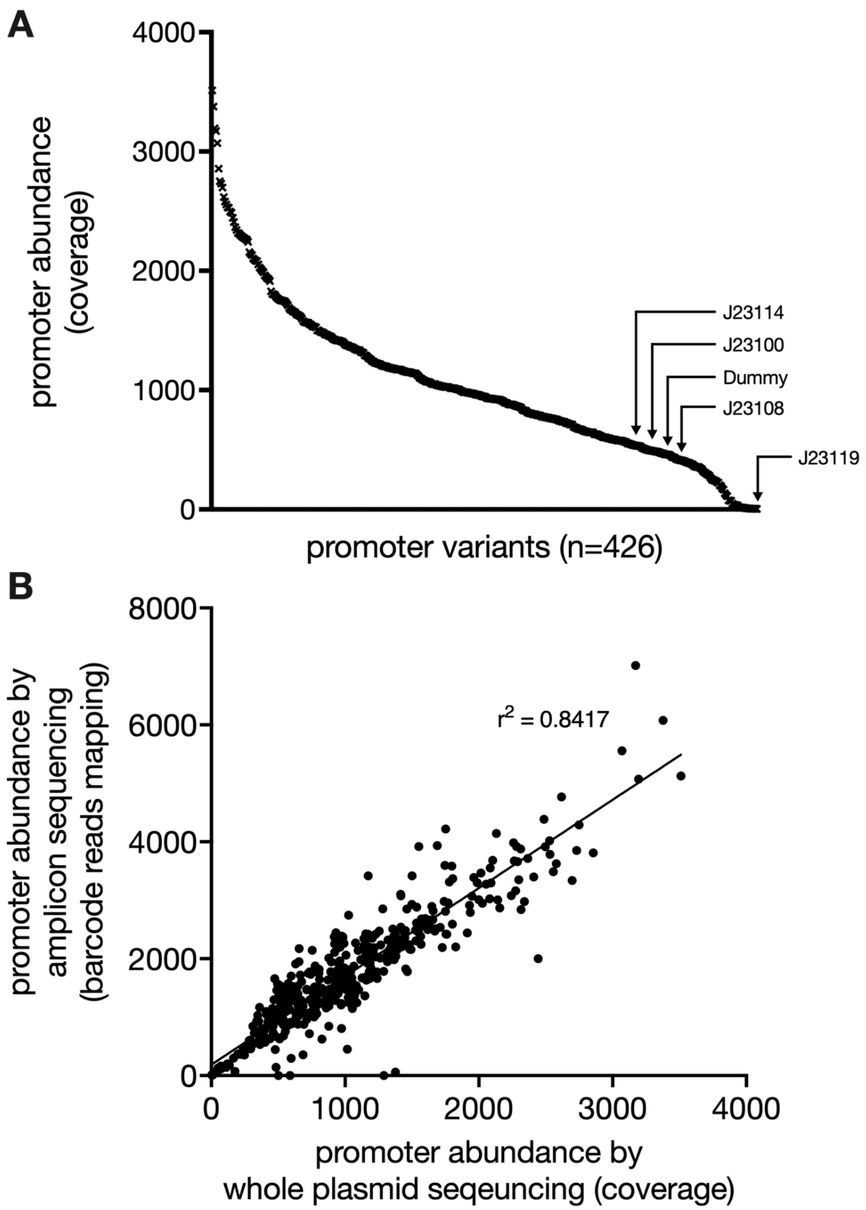


**Supplementary Figure 8. MACKEREL module Level 0 part composition.**
(A) Promoter variant abundance in MACKEREL module Level 0 part, as determined by whole plasmid sequencing. Promoter variants are sorted from highest to lowest abundance. Location of Anderson and dummy control promoter variants are indicated with arrows. (B) Correlation between promoter variant abundance in the MACKEREL module, as determined by two different next-generation sequencing methods: whole plasmid sequencing (X-axis) and amplicon-sequencing (Y-axis).

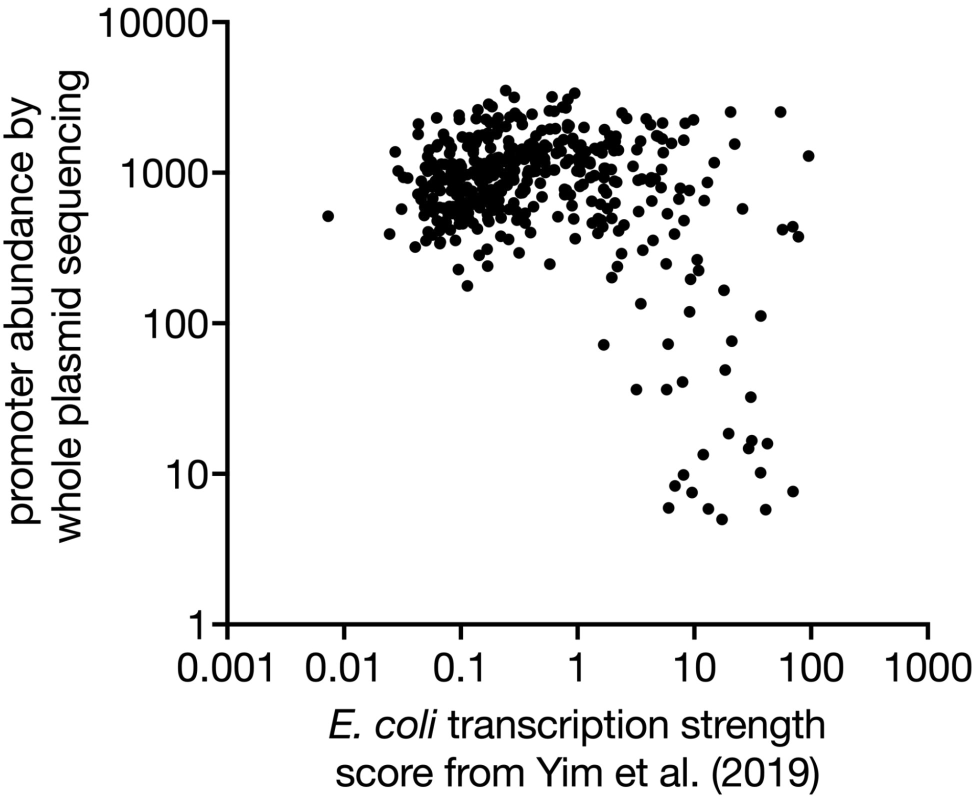


**Supplementary Figure 9. Correlation between MACKEREL module promoter abundance and transcription strength in *E. coli*.**
Promoter transcription strength scores for RS421 were obtained from Yim et al. (2019) and plotted against promoter abundance as determined by whole plasmid sequencing of the MACKEREL module. Note the Anderson library promoters (n=4) and the dummy promoter (n=1) present in the MACKEREL module are excluded from this analysis.

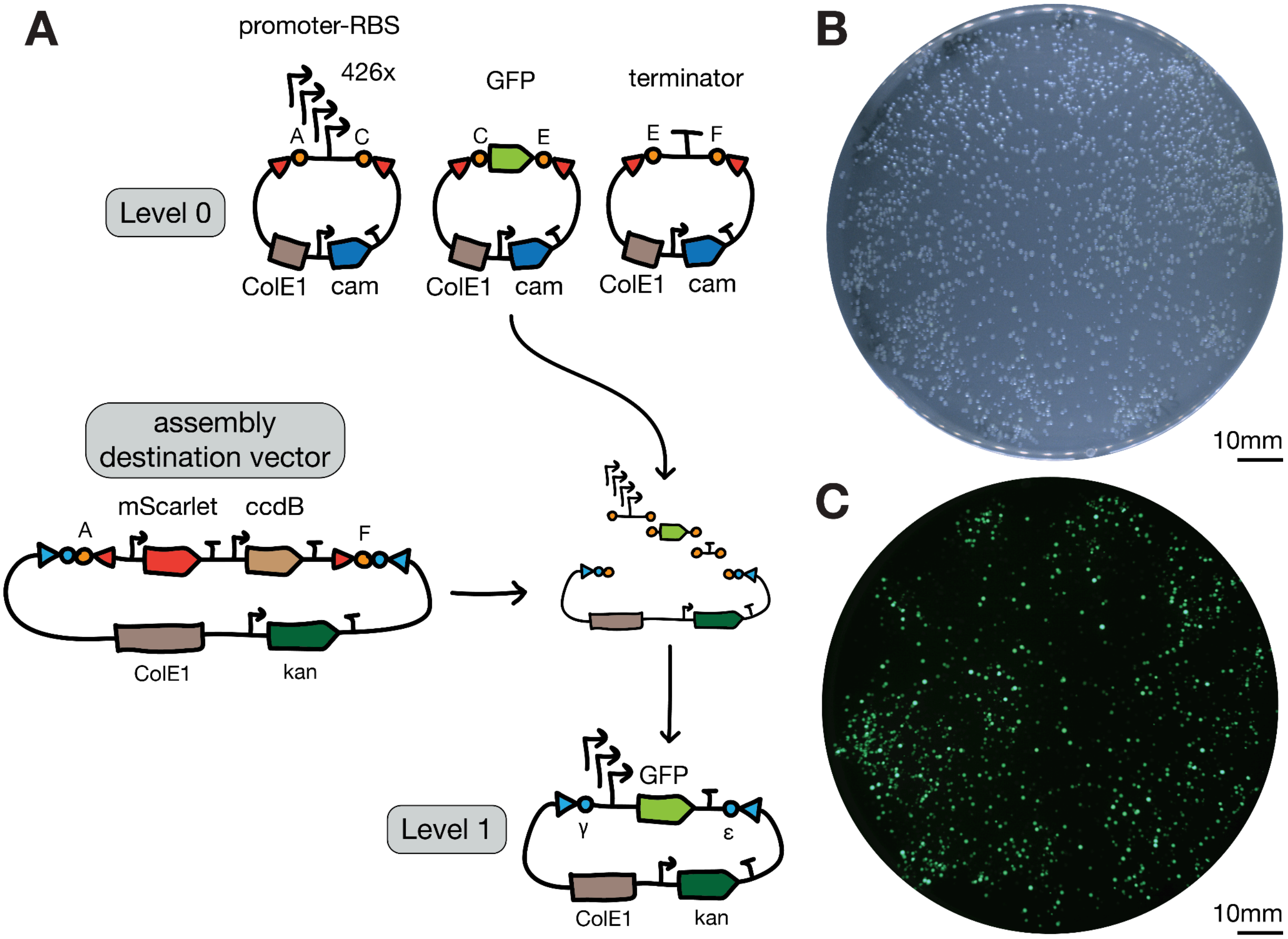


**Supplementary Figure 10. Level 1 assembly of the MACKEREL module with sfGFP.**
(A) Schematic of the Level 1 assembly procedure used to combine the MACKEREL module with sfGFP coding sequence. A non-standard assembly destination vector was used in this case where the standard sfGFP dropout part is replaced with an mScarlet-I-*ccdB* dropout part. The mScarlet-I cassette fulfills the same function as sfGFP in standard assembly destination vectors - enabling facile identification of re-ligated destination vector on transformation plates. The *ccdB* cassette is incorporated in an attempt to eliminate re-ligated destination vector from transformation plates. The *ccdB* gene encodes a toxin that is lethal to *E. coli*, preventing plasmid maintenance in the population. As a result, the destination vector itself must be maintained in specific *E. coli* strains that are resistant to the *ccdB* toxin, such as ccdB Survival 2 T1R *E. coli* (Invitrogen). (B) Brightfield and (C) green fluorescence images of a MACKEREL-sfGFP library *E. coli* transformation agar plate. A range of fluorescence intensities were observed across library transformants, consistent with the expected behavior of the MACKEREL module promoter variants.

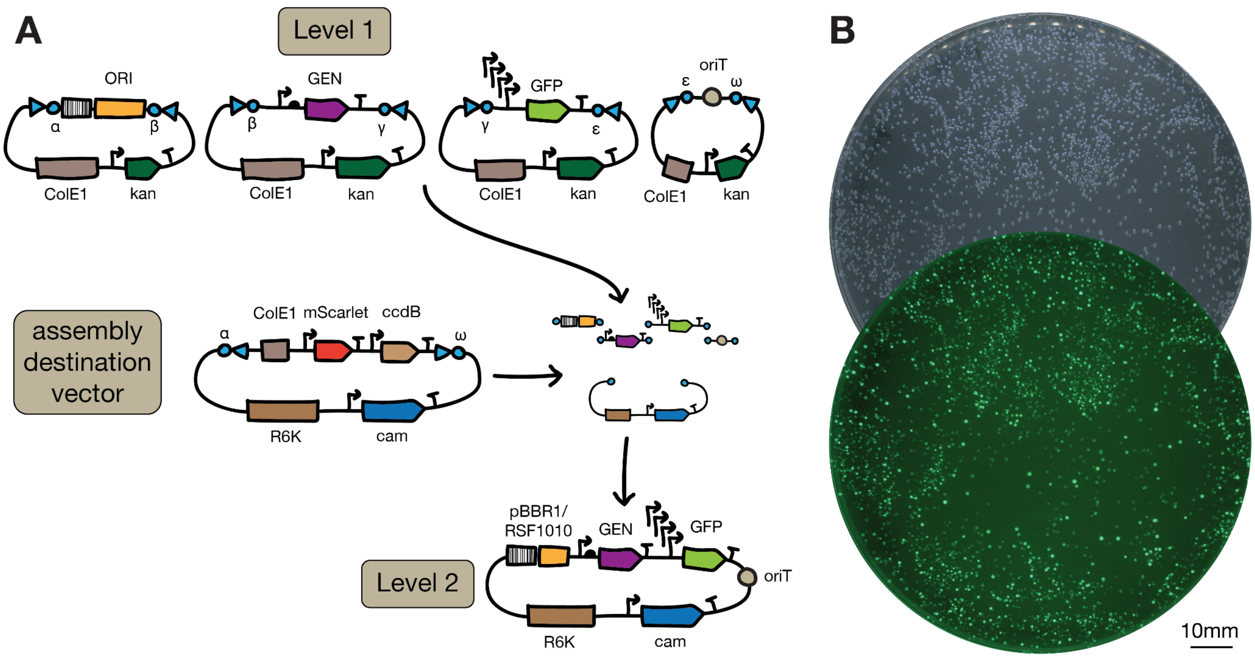


**Supplementary Figure 11. Level 2 assemblies of the MACKEREL module-sfGFP cassette with components required for non-model microbe transformation.**
(A) Schematic illustration of the Level 2 assembly procedure used to combine the MACKEREL module-sfGFP cassette with components required to transform non-model bacteria. The MACKEREL module-sfGFP Level 1 cassette was combined with: two plasmid ORIs demonstrated to replicate in the target strains (pBBR1 and RSF1010), a gentamicin resistance cassette demonstrated to function in the target strains, and the RK2/RP4 *oriT* sequence to enable delivery by conjugation. Once again, a non-standard assembly destination vector was used. As in Supplementary Figure 10, this destination vector possesses mScarlet-I and *ccdB* cassettes in the dropout part. In addition, the backbone possesses the conditional R6Kγ ORI for replication in *E. coli*. Since a *ccdB*-resistant, *pir*-plus *E. coli* strain was not immediately available, we also included a ColE1 ORI within the dropout part. The destination vector can therefore be maintained in ccdB Survival 2 T1R *E. coli* (Invitrogen), which is a *pir*-minus strain, *ccdB* resistant strain, and then transformed to *E. coli* pir-116 or *E. coli* BW29427, which are *pir*-plus but *ccdB* sensitive strains. Two final libraries were generated: pGL2_202 [RSF1010 ORI] and pGL2_204 [pBBR1 ORI]. (B) Brightfield and green fluorescence images of a pGL2_204 library transformation agar plate. A range of fluorescence intensities were observed across library transformants.

**A**

| Recipient species | # of library transconjugants (CFU) |
|---|---|
| *E. coli* | $2.1x10^7$ |
| *S. oneidensis* | $5.2x10^4$ |
| *S. putrefaciens* | $3.7x10^2$ |
| *H. alkaliphila* | $2.0x10^1$ |
| *H. neptunia* | $6.5x10^5$ |
| *H. elongata* | $1.6x10^2$ |
| *P. alcaliphila* | $5.3x10^2$ |

**B**

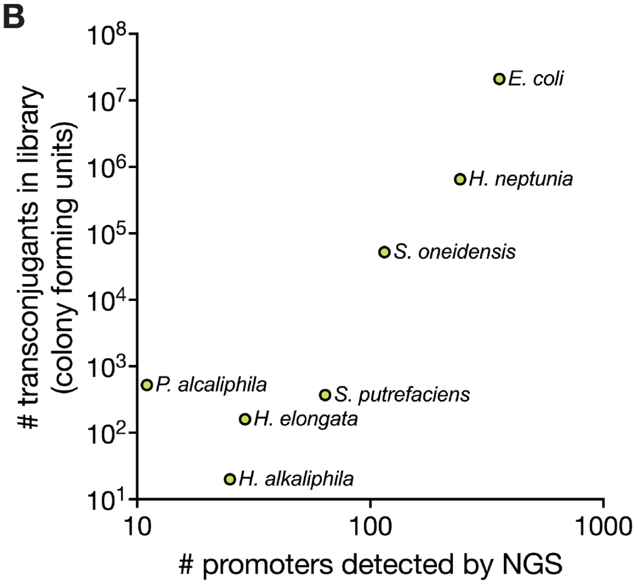


**Supplementary Figure 12. Characterization of MACKEREL module transconjugants.**
(A) Table showing the number of transconjugant colony forming units (CFUs) obtained from delivery of MACKEREL module constructs to recipient species. (B) XY scatter plot comparing the number of promoter library transconjugants obtained for each recipient species against the number of promoter variants detected in transconjugant libraries by NGS.

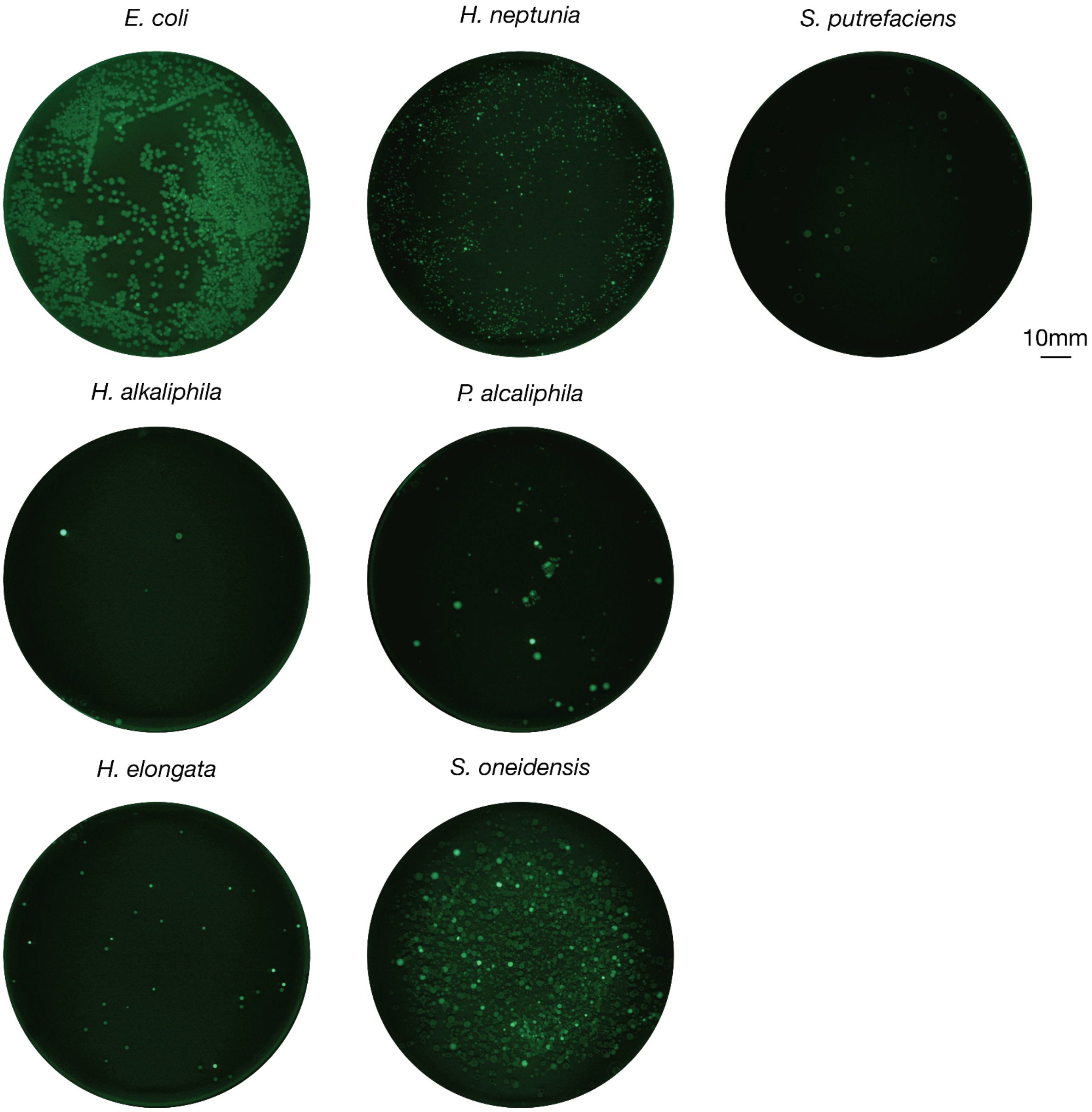


**Supplementary Figure 13. Fluorescence imaging of MACKEREL module transconjugant plates.**

Green fluorescence images of transconjugants obtained following MACKEREL module delivery to a seven target species. A range of fluorescence intensities were observed across transconjugants.
